# A new *Opn4cre* recombinase mouse line to target intrinsically photosensitive retinal ganglion cells (ipRGCs)

**DOI:** 10.1101/2024.04.16.589750

**Authors:** Brannen Dyer, Sue O. Yu, R. Lane Brown, Richard A. Lang, Shane P. D’Souza

## Abstract

Intrinsically photosensitive retinal ganglion cells (ipRGCs) play a crucial role in several physiological light responses. In this study we generate a new *Opn4^cre^* knock-in allele (*Opn4^cre(DSO)^*), in which cre is placed immediately downstream of the *Opn4* start codon. This approach aims to faithfully reproduce endogenous *Opn4* expression and improve compatibility with widely used reporters. We evaluated the efficacy and sensitivity of *Opn4^cre(DSO^*^)^ for labeling in retina and brain, and provide an in-depth comparison with the extensively utilized *Opn4^cre(Saha^*^)^ line. Through this characterization, *Opn4^cre(DSO)^* demonstrated higher specificity in labeling ipRGCs, with minimal recombination escape. Leveraging a combination of electrophysiological, molecular, and morphological analyses, we confirmed its sensitivity in detecting all ipRGC types (M1-M6). Using this new tool, we describe the topographical distributions of ipRGC types across the retinal landscape, uncovering distinct ventronasal biases for M5 and M6 types, setting them apart from their M1-M4 counterparts. In the brain, we find vastly different labeling patterns between lines, with *Opn4^cre(DSO)^* only labeling ipRGC axonal projections to their targets. The combination of off-target effects of *Opn4^cre(Saha)^* across the retina and brain, coupled with diminished efficiencies of both *Cre* lines when coupled to less sensitive reporters, underscores the need for careful consideration in experimental design and validation with any *Opn4^cre^*driver. Overall, the *Opn4^cre(DSO^*^)^ mouse line represents an improved tool for studying ipRGC function and distribution, offering a means to selectively target these cells to study light-regulated behaviors and physiology.

## INTRODUCTION

Intrinsically photosensitive retinal ganglion cells (ipRGCs) represent a diverse class of retinal output neuron with 6 types (M1-M6) classified by melanopsin content, dendritic stratification, gene expression, and electrophysiological response characteristics^1,2^. A key shared feature of M1-M6 is their expression of melanopsin (encoded by *Opn4*) and thus their ability to autonomously detect light that causes the cell to depolarize^1^. While some types, like the M1-M3 ipRGCs, display strong photocurrents and express sufficient photopigment to be detected by typical antibody staining approaches^3^, M4-M6 have limited melanopsin immunoreactivity and correspondingly weak photocurrents^4^. Functionally, ipRGCs convey photic information to diverse brain regions, serving as a major conduit for the effects of light on circadian entrainment^5–7^, the pupillary light response^8,9^, mood^10,11^, sleep^12^, learning^10,11^, and pattern vision^13,14^.

The discovery of ipRGCs and their functions has been driven by the development of genetic tools that express recombinases (*Cre*)^14^, toxins (DTA, attnDTA)^9,15^, or reporter genes (GFP, tauLacZ, tdTomato) from the *Opn4* locus^6,15,16^. However, due to variable expression of melanopsin within each ipRGC type, these lines vary considerably in their specificity and selectivity. For example, *Opn4^tauLacZ^* primarily labels M1 ipRGCs due to their high expression of melanopsin^6^, while the *Opn4-GFP* transgenic line tends to label types with higher melanopsin content^16^. The most sensitive of these methods, labeling all ipRGC types, utilizes *Cre* recombinase expressed from the *Opn4* locus (*Opn4^cre(Saha)^*) in conjunction with a *Cre*-dependent reporter, and has provided insight into ipRGCs diversity^14^. Additionally, when coupled to *Cre*-dependent loss-of-function alleles (floxed genes) or *Cre*-dependent intersectional ablation lines (*Pou4f2^zDTA^*), the use of *Opn4^cre(Saha)^* has uncovered several type-specific and functional roles for ipRGCs in light-regulated behavior and physiology^11,17–19^

When crossed with sensitive *Cre*-dependent reporters (*Ai9* & *Ai14*)^20^, *Opn4^cre(Saha)^*produces recombination in mostly non-ipRGC cell types within the retina and across the brain^21^. Conversely, when crossed to less sensitive reporters such as *Z/EG*, in addition to recombination in outer retinal photoreceptors^14^, it has been reported that ∼30% of melanopsin immunoreactive cells are not labeled^21,22^, suggesting the specificity and sensitivity of this *Cre* recombinase line are highly reporter-dependent. One explanation for this may be the design of the *Opn4^cre^*^(^*^Saha^*^)^ line (Figure 1 bottom). In this line, *Cre* recombinase was knocked into the *Opn4* locus, replacing most of the gene, and a human beta globin intron was incorporated upstream of the *Cre* coding sequence. This was likely motivated by the desire for robust expression and efficient deletion of floxed alleles in weakly-expressing cells such as M4-M6 ipRGCs^4^. However, given that only a few molecules of Cre-recombinase may be sufficient to achieve recombination in the genome^23^, this additional regulatory element could drive off-target effects depending on the type of reporter used for tagging or accessibility of the floxed gene being studied.

**Figure 1.**
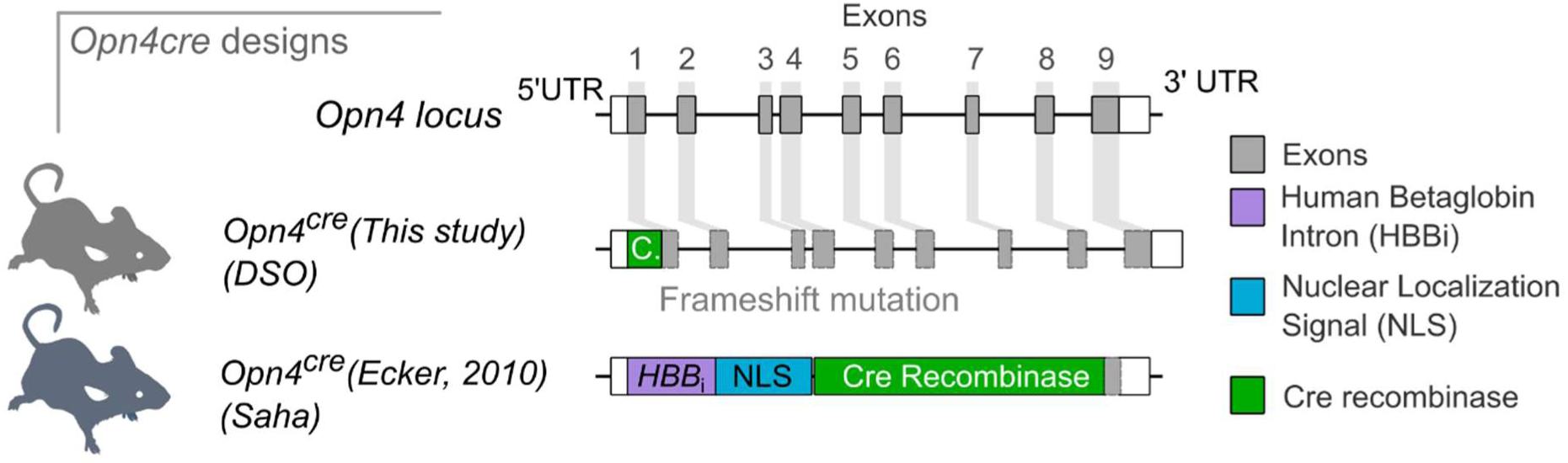
The design of *Opn4^cre^* alleles. *Opn4* gene structure in the mouse (top). Design of the *Opn4^cre(DSO^*^)^ line by CRISPR-Cas9 mediated knock-in of *Cre* recombinase, introducing a frameshift mutation in *Opn4* but retaining gene components and structure (middle). Design of the *Opn4^cre(Saha^*^)^ line (lower). In this allele, the majority of the *Opn4* gene is deleted and replaced with a *Cre* cassette comprising an upstream human beta globin intron (HBBi) and a nuclear localization signal (NLS). UTR = untranslated region.

In this study we describe and characterize a new knock-in *Opn4^cre^*line (termed *Opn4^cre(DSO)^*) where *Cre* is knocked-in immediately downstream of the *Opn4* start codon, leading to disruption of the gene but preservation of the downstream gene structure. We compared this line to *Opn4^cre(Saha)^* by crossing to widely used reporters and assessed their labeling efficiency within the retina and brain. We find that *Opn4^cre(DSO)^* specifically labels ipRGCs within the retina, and has low recombination escape for M1-M3 ipRGCs. Additionally, using a combination of electrophysiology, molecular markers, and morphology, we find that this line is sensitive, capable of detecting all ipRGC types previously described (M1-6). Using this tool, we describe the topographical distribution of these types across the retina from adult mice, showing the M5 and M6 ipRGCs are spatially biased towards the central-ventronasal retina, unlike their M1-M4 counterparts^24–26^. In addition, we catalogue the labeled cells within the brain using both *Cre* lines. We find that *Opn4^cre(Saha)^* labels multiple brain regions at high density while *Opn4^cre(DSO^*^)^, shows low density sporadic activity with no specific pattern. Finally, using a *Cre*-dependent reporter with larger inter-*loxP* distances (*mTmG*) but identical genomic accessibility to *Ai14*, we find that both *Opn4^cre^* lines have reduced efficiency of labeling ipRGCs. These data define the characteristics of *Opn4^cre(Saha)^*and *Opn4^cre(DSO)^* and inform decisions about their use.

## RESULTS

### *Opn4^cre(DSO)^* mouse line generation

To generate a mouse line with specific expression of *Cre* from the *Opn4* locus, we employed a CRISPR-Cas9 targeting approach in C57BL/6J embryos (see Methods for detail) that introduced a cut upstream of the *Opn4* start codon. A co-injected long single stranded DNA (ssDNA) containing the coding sequence of *Cre*-recombinase, the SV40 polyA element, and homology arms to the 5’ UTR and exon 1 served as a donor. This design places expression of *Cre* under the control of the endogenous *Opn4* promoter. In this design, the downstream coding sequence is disrupted by a frameshift, but the gene elements (exons 1-9, introns) remain unchanged (Figure 1). Using this approach, we generated a single founder male that was used to propagate the *Opn4^cre(DSO)^* genetic line.

### *Opn4^cre^* ipRGC efficiency, sensitivity, and off-target labeling when crossed to a sensitive reporter

To compare the targeting efficiencies of both *Opn4^cre^*lines, we crossed them to *Ai14* mice^20^, one of the most widely used *Cre*-dependent reporters. This line harbors a *loxP*-3xSTOP-polyA-*loxP* cassette upstream of a tdTomato fluorescent reporter located in the highly accessible *ROSA26* locus. Upon *Cre*-mediated excision of the stop cassette, strong tdTomato expression is driven by an upstream CAG promoter. From these crosses, we analyzed expression patterns of tdTomato within the inner retina of wholemount samples while simultaneously immunolabeling ipRGCs for Opn4 (melanopsin). When crossed to *Ai14*, the newly developed *Opn4^cre(DSO)^*line expressed tdTomato primarily in the ganglion cell layer (GCL; 3264 ± 170 cells, *n* = 8) and within the inner nuclear layer (INL; 800 ± 47 cells, n = 4), where *Opn4*-expressing ipRGCs reside (Fig 2a). In comparison, tdTomato+ cells were far more abundant in the *Opn4^cre(Saha^*^)^*; Ai14* retina, with 10,737 ± 148 cells in the GCL (*n =* 4), and 1795 ± 186 cells within the INL (*n =* 2) (Fig 2c). In contrast, it was previously reported that only ∼2000 cells per retina were labeled in the *Opn4^cre(Saha)^*; *Z/EG* mouse line^14,21^.

**Figure 2.**
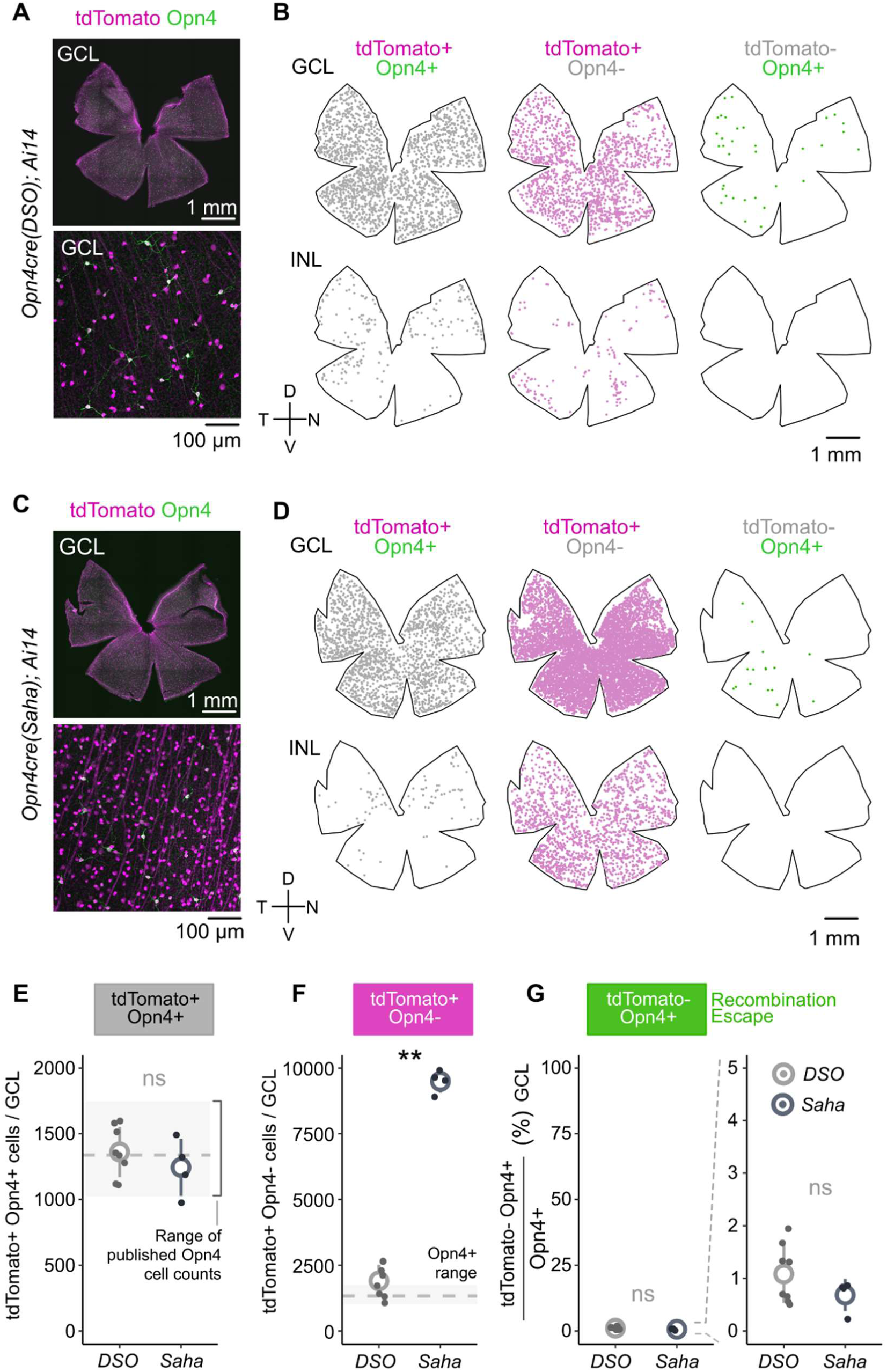
ipRGC labeling efficiency within the inner retina of each *Opn4cre* line. (A) Representative flat-mount retina from *Opn4^cre(DSO)^*; *Ai14* mice. (B) Reconstruction of flat-mount retina in (A) colored by combination of tdTomato expression (*Ai14,* magenta) and Opn4 co-expression (green) in both the GCL (top row) and INL (bottom row). (C-D) Representative flat-mount and reconstructed retina from *Opn4^cre(Saha)^*; *Ai14* presented across the GCL and INL as in (A-B). (E-G) Analysis of *Cre*-driven recombination across both *Cre* lines crossed to *Ai14*. (E) Absolute cell counts of tdTomato+ Opn4+ cells in the GCL (*DSO_n_* = 8; *Saha_n_* = 4 animals) with grey box depicting min and max range of published, dashed line representing mean of published counts. (F) Same as E but comparing tdTomato+ Opn4-cells in the GCL, grey box and line are mirrored from (E) (\*\**p <* 0.01, Wilcoxon signed-rank test). (G) Percentage of Opn4+ cells that are tdTomato-in the GCL. GCL = ganglion cell layer; INL = inner nuclear layer. D = dorsal, N = nasal, V = ventral, T = temporal. Small dots = individual samples (*n*), large circles = group means, error bars = standard deviation.

To assess the efficiency of *Cre*-mediated recombination, we categorized cells into 3 groups following tdTomato and Opn4 labeling. **(1)** Those that have recombined *Ai14* to express tdTomato and express Opn4, reflecting M1-M3 types (tdTomato+ Opn4+), **(2)** cells that have recombined *Ai14* but lack Opn4-immunoreactivity (tdTomato+ Opn4-), comprising the M4-M6 types and/or off-target labeling, and, **(3)** cells that express Opn4 protein, but had not recombined *Ai14* (tdTomato-Opn4+) (Fig 2b, d). Across the GCL, we found a large disparity in the tdTomato+ Opn4-population between the two *Opn4^cre^* lines (*DSO*: 1904 ± 217, *Saha*: 9494 ± 215 cells) (Fig 2f), but similar numbers of tdTomato+ Opn4+ cells (*DSO*: 1360 ± 68, *Saha*: 1243 ± 109 cells) and cells that evaded recombination (*DSO*: 14 ± 2, *Saha*: 9 ± 2 cells). In both lines, double positive cells were within the range of previously published counts for Opn4-immunoreactive ipRGCs (Fig 2e)^21,25,27–30^, with ∼1% or less that escaped (Fig 2g). These data argue that both lines, even with differing strategies for driving *Cre* expression, are efficient at recombining a sensitive reporter in Opn4+ cells with minimal escape rates.

Given the nearly 5-fold difference in tdTomato+ Opn4-cells between the *Opn4^cre^* lines, we sought to better understand whether the abundance in *Opn4^cre(Saha)^* cells reflected higher sensitivity or off-target labeling. To this end, we assessed tdTomato+ cell distributions in retinal cross-sections, which allowed for a more detailed, layer-specific analysis of labeled cells (S. Fig 1a-b). As observed in wholemount analysis, *Opn4^cre(DSO)^* retina contained fewer tdTomato+ cells in both the GCL and INL. We also found that *Opn4^cre(DSO)^* lacked any cell labeling in the outer nuclear layer (ONL), where rod and cone photoreceptors reside (S. Fig 1a-b). By contrast, *Opn4^cre(Saha^*^)^ produced abundant cell labeling in the ONL, the majority of labeled cells within the retina (312.6 ± 44.3 cells mm^-2^). Recombination in the outer retina is not an artefact of *Ai14*, as the *Opn4^cre(Saha^*^)^; *Z/EG* line also shows abundant labeling in outer retinal photoreceptors^14^. Given that there is limited data to suggest rod and cone photoreceptors express melanopsin, and there is no evidence of outer photoreceptor labeling in the *Opn4^cre(DSO^*^)^ line, it is likely this reflects line-specific off-target labeling.

### Molecular fingerprinting of *Opn4^cre^* labeled cells

Melanopsin expression is thought to be restricted to RGCs, which are located primarily in the GCL with a small fraction displaced in the INL^5,22,31^. Given that 11.6% (*Saha*) to 41.6% (*DSO*) of tdTomato cells express Opn4, we set out to determine whether the remaining inner retinal cells represent RGCs or other cell types. To this end we stained retina for Rbpms, a selective marker of RGCs^32^, and determined, in both *Opn4^cre^* lines, the extent of colocalization with tdTomato. In the *Opn4^cre(DSO^*^)^ line, nearly all tdTomato+ cells were RGCs across the GCL (48/48 cells, S Fig 2a-b) and INL (3/4 cells, S Fig 2a-b). By contrast, in the *Opn4^cre(Saha)^* line, 73.8% of tdTomato+ cells in the GCL (79/107 cells) and 5.6% in the INL (1/18 cells) appear to be RGCs (S. Fig 2a-b). Considering that >10,000 cells were labeled within the GCL of the *Opn4^cre(Saha)^* line (Fig. 2), approximately ∼2800 are likely to be non-RGCs.

Due to low density of cells in cross-section we also performed Rbpms immunolabeling in flat-mounts from *Opn4cre^(DSO)^; Ai14* mice. Wholemount analysis of tdTomato/Rbpms overlap across the inner retina (Fig 3a-d) revealed that nearly all cells across animals were RGCs (tdTomato+ Rbpms+ = 99.6 ± 0.37 %, *n* = 4). Spatially, and consistent with previous reports, INL-localized ipRGCs appeared to be enriched in the periphery of the temporal retina^28^ and the ciliary marginal zone of the nasal retina^30^ (Fig 3d, INL tdTomato+ Rbpms+ = 91.1 ± 8.48 %, *n* = 4). With the exception of one example of sporadic and spatially restricted recombination within the INL (Fig 3b, d; sample #3, *n =* 1/18 animals assessed), these results suggest this *Opn4^cre(DSO)^* specifically marks RGCs, consistent with the known expression patterns of *Opn4* and spatial distribution of the ipRGC population.

**Figure 3.**
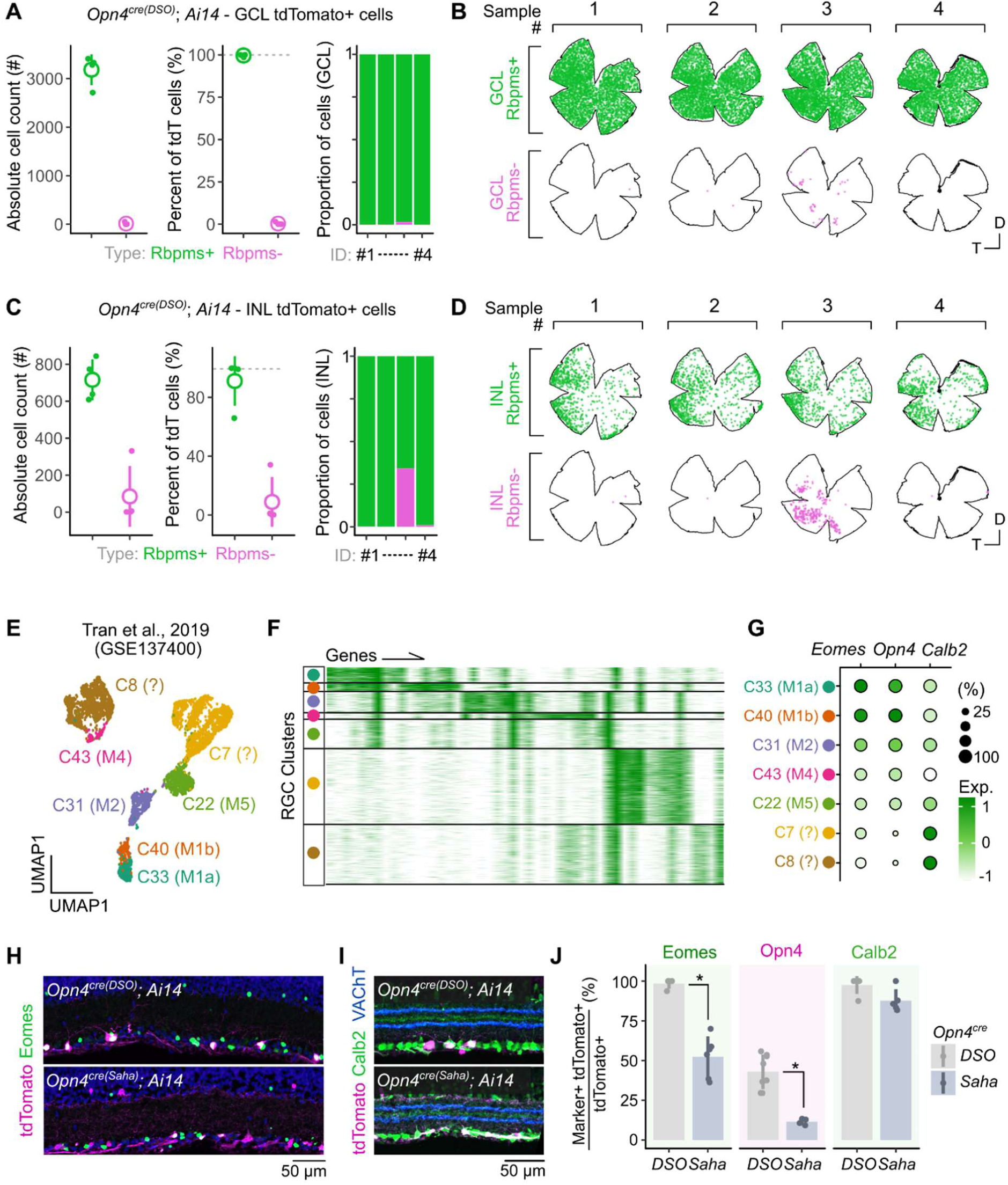
Molecular characterization of cells labeled in the *Opn4^cre(DSO)^* mouse line. (A) Colocalization analysis of Rbpms in tdTomato+ cells across the GCL in the *Opn4^cre(DSO^*^)^ line as absolute counts (left), percent of tdTomato+ cells (middle) and individual sample proportion plots (right) (*n =* 4 animals). (B) Representative spatial analysis of Rbpms and tdTomato colocalized cells in the GCL. (C) Same samples and analysis as in (A) but within the INL. (D) Same samples and analysis as in (B) but within the INL. (D-E) Reanalysis of Tran & Shekhar et al., 2019 to identify shared and unique genes between ipRGC (C22-C43) and putatively-ipRGC (C7, C8) clusters. (D) 2D UMAP embedding of clusters from Tran & Shekhar et al., 2019. (E) Heatmap depiction of significantly differentially expressed genes (DEGs) between each cluster, ordered by cells as rows and genes as columns. (F) Dotplot of shared ipRGC genes *Eomes, Opn4, and Calb2*. Expression scale bar is identical between (E) and (F). (G-I) Validation and comparison of marker expression in tdTomato+ cells from both *Cre* lines crossed to *Ai14*. (G-H) Representative confocal images of Eomes staining (G) and Calretinin staining (H). (I) Comparison of tdTomato+ colocalization with different markers between both *Cre* lines (Eomes*_DSO vs Saha_* _*_*p* < 0.05; Opn4*_DSO vs Saha_* \**p* < 0.05; Calb2*_DSO vs Saha_ p* > 0.05; Wilcoxon signed-rank test with *fdr* correction). Bar height = mean, error bars = standard deviation.

We sought more specific markers of ipRGCs to determine the specificity of each *Cre* line and reanalyzed an existing single cell RNA-Seq atlas of RGC types in the adult mouse^33^ (Fig 3e-f). In this analysis, we included clusters C33 (M1a), C40 (M1b), C31 (M2), C43 (M4), C22 (M5), C7 (Novel), and C8 (Novel). These RGC clusters share expression of the transcription factor *Eomes* which is expressed in all ipRGC types and displaced amacrine cells^34^. The novel clusters, C7 and C8, lack *Opn4* expression and though considered *Eomes*+ RGCs, express the least amount of *Eomes* (Fig 3g). Thus, it is currently unclear if they represent ipRGCs (M1-M6) or whether they represent transcriptionally related non-ipRGC types^34^. Additionally, *Calb2* which encodes calretinin, is expressed broadly by most RGCs and types of amacrine cells^35^. We labeled retina from *Opn4^cre(DSO)^; Ai14* and *Opn4^cre(Saha^*^)^; *Ai14* against Eomes (Fig 3h) and calretinin (*Calb2,* Fig 3i). Almost all labeled cells in the *Opn4^cre(DSO^*^)^ line expressed Eomes (98.2 ± 1.8 % Eomes+). This compared with 52.4 ± 5.2 % overlap in the *Opn4^cre(Saha^*^)^ line (Fig 3j). Calretinin co-labeling did not vary between different lines (Fig 3i). Together, these data suggest that when crossed to *Ai14*, the *Opn4^cre(DSO^*^)^ line primarily labels ipRGCs, while *Opn4^cre(Saha)^* labeling also includes a distribution of different RGC and amacrine types in the inner retina.

Consistently, analysis of tdTomato+ dendrite stratification in the inner plexiform layer (IPL) revealed spatially restricted signal in the *Opn4^cre(DSO)^*line that was highest at the inner (S5) and outer (S1) boundaries of the IPL, where ipRGC dendrites stratify (S. Fig. 3a-c)^36^. In contrast, dendritic signals were much more diffuse in the *Opn4^cre(Saha)^* line, consistent with previous reports, with a notable peak at the outer boundary of the IPL (S1) (S. Fig. 3d-f). Taken together, the *Opn4^cre(DSO^*^)^ line specifically labels ipRGCs within the inner retina.

### Electrophysiological profiling of RGCs in the *Opn4^cre(DSO)^* line

A hallmark of ipRGCs is their light-evoked intrinsic photocurrent in the absence of synaptic input from the rod and cone pathways^5,16^. Though intrinsic responses to light vary in both amplitude and kinetics^4^, all ipRGCs respond to a full-field flash of light with an inward current, even if their melanopsin content is extremely low^4^. Given that the *Opn4^cre(DSO^*^)^ line primarily labels ipRGCs based on molecular analyses, we assessed their intrinsic responses to a full-field flash of 480 nm light (Fig 4a-b). Whole-cell voltage-clamp recordings revealed photocurrents in 95.2% of cells (20/21 tdTomato+) patched in the *Opn4^cre(DSO^*^)^ line crossed to *Ai9*. Peak inward photocurrents varied from 10 to 500 pA, consistent with intrinsic responses from M1-M6 ipRGCs^3,13,37–40^. Due to the heterogeneity of responses, we clustered photocurrents based on their temporal properties. We identified 3 photo-responsive clusters (C1,2,4) and a cluster that contained the non-responsive cell, which appeared to have current fluctuations prior to stimulus onset (C3)(Fig 4c). Cluster C1 represents cells with the strongest photocurrents (average peak inward current = 367.8 pA), while clusters C2 & C4 had substantially weaker intrinsic responses to light (peak inward current, C2 = 36.6 pA; C4 = 11.4 pA). A subset of cells was dye-filled with neurobiotin following recordings, allowing us to reconstruct ipRGC morphologies and match them to their intrinsic responses (S. Fig. 4A-C). A cell that mapped to cluster C1 (peak inward current of 339.8 pA), had sparsely branched dendrites stratified deep in the IPL, consistent with the response and morphology of M1s (Fig 4d)^3,5,8,27^. Cells that mapped to C2 (Fig 4e) and C4 (Fig 4f) had weaker photocurrents, and had more branched monostratified dendrites localized to the inner boundary of the IPL (S5). These features suggest these cells may represent the M5^26,39,40^ (C2), and M4^13,24,26^ (C4) type. Thus, we find a wide range of intrinsic photocurrents and morphological types in the *Opn4^cre(DSO^*^)^ line, consistent with existing data describing ipRGCs.

**Figure 4.**
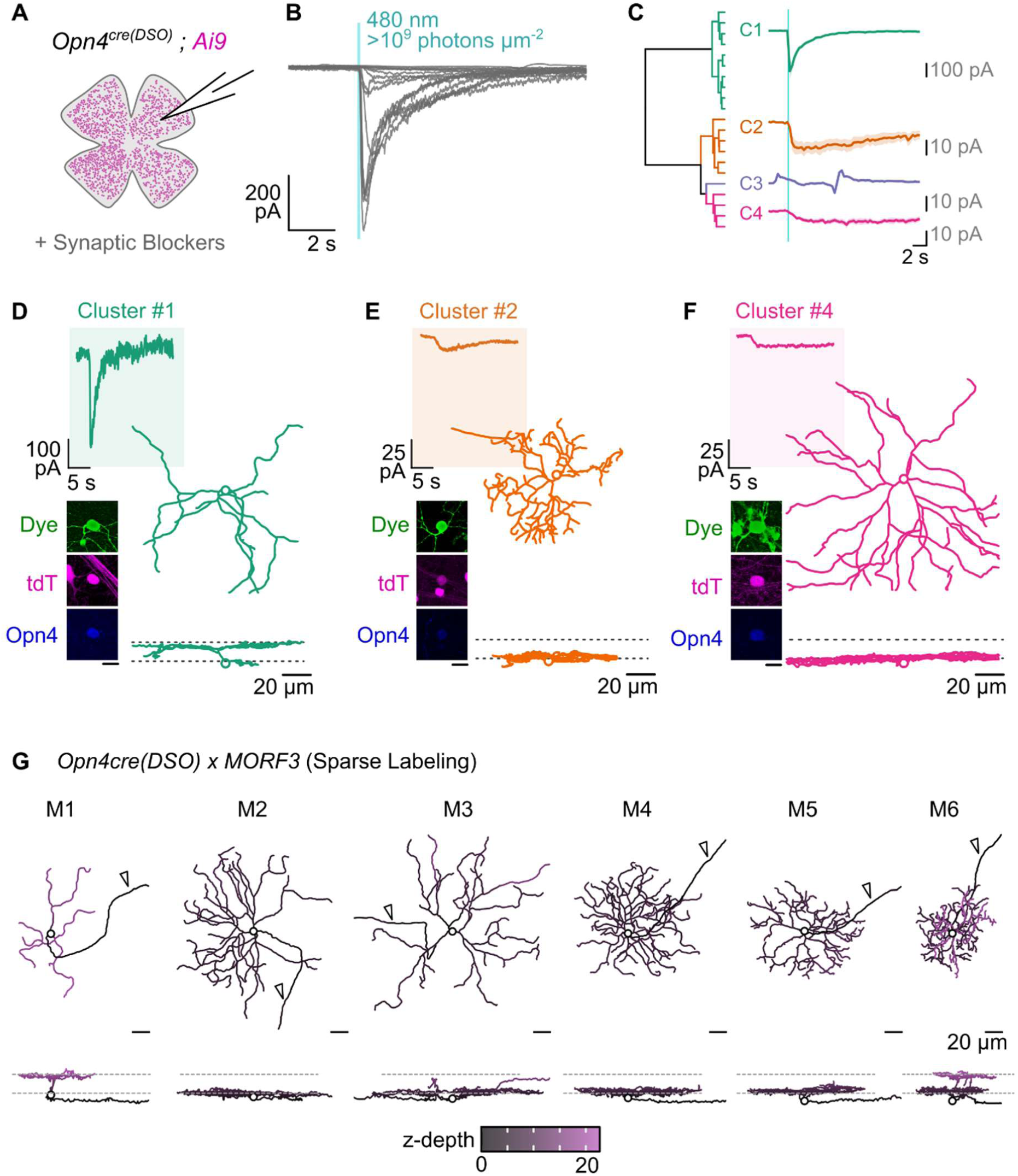
Electrophysiological and morphological assessment of ipRGCs in the *Opn4^cre(DSO^*^)^ line. (A) Schematic depicting whole-cell recording of *Opn4^cre(DSO)^; Ai9* cells from the retina. (B) Current traces of all recorded cells held at -60mV and given a 100 ms full-field flash of > 9 log photons um^-2^ sec^-1^, cyan region = light pulse. (C) Hierarchical clustering dendrogram and cluster averaged responses to light, bold line = mean response, shaded region = standard error. (D-F) Representative responses, confocal images, and 3-dimensional morphological reconstructions of cells from cluster 2 (D), cluster 3 (E), and cluster 5 (F). (G) Representative 3-dimensional reconstructions of ipRGC types from *Opn4^cre(DSO)^; MORF3* retina (*n* = 5) with XY profile (top) and XZ profile (bottom) with segments color-coded by z-depth. S = seconds. Arrowhead = axon.

### Morphological diversity of ipRGCs in the *Opn4^cre(DSO)^ line*

The finding that majority of cells in the *Opn4^cre(DSO^*^)^ line lack melanopsin-immunoreactivity but still represent ipRGCs due to their expression of Rbpms and Eomes (Fig 2), coupled with the diverse electrophysiological responses and morphologies, prompted a more robust characterization of ipRGC diversity. M1-M6 ipRGCs are readily distinguished morphologically as they vary in their dendritic stratification, branching, and soma size^2,4^.

However, to unequivocally assign cells to specific types requires reconstructing their full dendritic arbors with minimal overlap from other cells in close proximity. Thus, to genetically and sparsely label ipRGCs, we crossed the *Opn4^cre(DSO^*^)^ line to *MORF3* mice^41^. This *Cre*-dependent reporter utilizes mononucleotide repeat frameshift (MORF) as a switch for sparse labeling. For cells to express smV5 (spaghetti monster V5) epitopes, two events must take place: cells must express *Cre* to excise a *loxP*-flanked STOP cassette (as in *Ai14*) and must have undergone a spontaneous frameshift mutation in the polycytosine (C22) MORF switch upstream of V5. In the cells that these events do occur, V5 is detectable via antibody labeling and can be coupled with other markers^41^. Our prior work using this method estimated ∼3% of ipRGCs were labeled in this approach^42^ and thus we implemented a similar strategy to study the range of ipRGC types targeted by our line.

We surveyed 356 cells (*n =* 5 animals) and performed 3D reconstruction on a subset of neurons with representative morphologies (Fig 4g). We found that cells labeled in the *Opn4^cre(DSO^*^)^ line match the morphological characteristics of all known ipRGC types.

- M1 types that laminate their dendrites in S1 (outer boundary of the IPL) had sparsely branching monostratified dendritic trees^3,5,6,8,27^.
- M2 types had more branched dendritic trees than the M1s that laminated in S5, close to their somas^3,27^.
- M3 types were rare, but the few we found had similar branching densities to the M2 but stratified its dendrites in both S1 and S5^27,37^.
- M4 types, like M2s, were monostratified in S5 but had the largest somas of cells in this line, were more branched, and expressed the highest V5 content based on immunofluorescence^13,26,38^.
- M5 types were monostratified in S5 with dense, small dendritic fields with small somas^14,26,39^.
- M6, like the M5, had small dendritic fields and somas but were bistratified (S1 & S5), occasionally containing recurrent dendrites that returned to S5 after branching in S1^40^.

### Topographical distributions of ipRGC types revealed by the *Opn4^cre(DSO^*^)^ line

Nonuniformities in the spatial distribution of retinal ganglion cell types serve to enhance encoding of specific visual features^24,43^. Examples include the lateral positioning of ipsilaterally projecting RGCs that allow for binocularity, to the heightened RGC densities of the primate fovea and area centralis of carnivores that enhance high-resolution vision^43,44^. Such spatial anisotropies are not limited to image-forming RGC types, as many ipRGCs display spatial enrichments. The pSON-projecting M1 subtype is nearly exclusively located in the dorsal retina^45^, while the remainder of the M1 population is evenly distributed across the dorso-ventral axis but is enriched temporally^30,45^. Similarly, the M2 type is thought to be enriched dorsally^25^, while the M4 (also known as Alpha ON-sustained RGC) has the highest density in the temporal retina^24,26^. However, it remains unclear if the other ipRGCs types display nonuniform distributions in retinal space. Previous work that first described the M6 type used the *Cdh3-GFP* mouse line that shows dynamic expression over development and strain^40,46^, and thus cannot provide much insight into spatial distributions of the M6 ipRGC. Given that the *Opn4^cre(DSO^*^)^ line specifically labels ipRGCs, including the weakly melanopsin-expressing M5 and M6 types (Fig. 4), we set out to understand whether each ipRGC type is nonuniformly distributed through the retina.

We first used an approach that coarsely defined ipRGC types in *Opn4^cre(DSO^*^)^; *Ai14* retina (Fig 5a-c). Immunostaining of tdTomato+ cells for melanopsin (Opn4) reveals the M1-M3 types, while reactivity to SMI32 (an antibody generated against the non-phosphorylated form of neurofilament heavy chain) selectively marks M4s^13,26^. The remaining cells that are defined by lack of melanopsin or SMI32 staining are considered the M5 and M6 type. As a quality control step, we surveyed the entire retina (GCL, INL, and ONL) for any off-target labeling which might lead to a false positive M5/6 assignment. We then assessed the spatial positions of each soma for three broad classes (M1-M3, M4, and M5-M6), with a representative flat mount highlighted in Fig 5c. To eliminate the spatial distortions introduced by relaxing cuts on the retina made during preparation, we reconstructed retina into a standard, spherical retinal space. Beyond limiting cut-site distortions, this approach allows for a registered space to compare spatial distributions of several ipRGC and photoreceptor populations.

**Figure 5.**
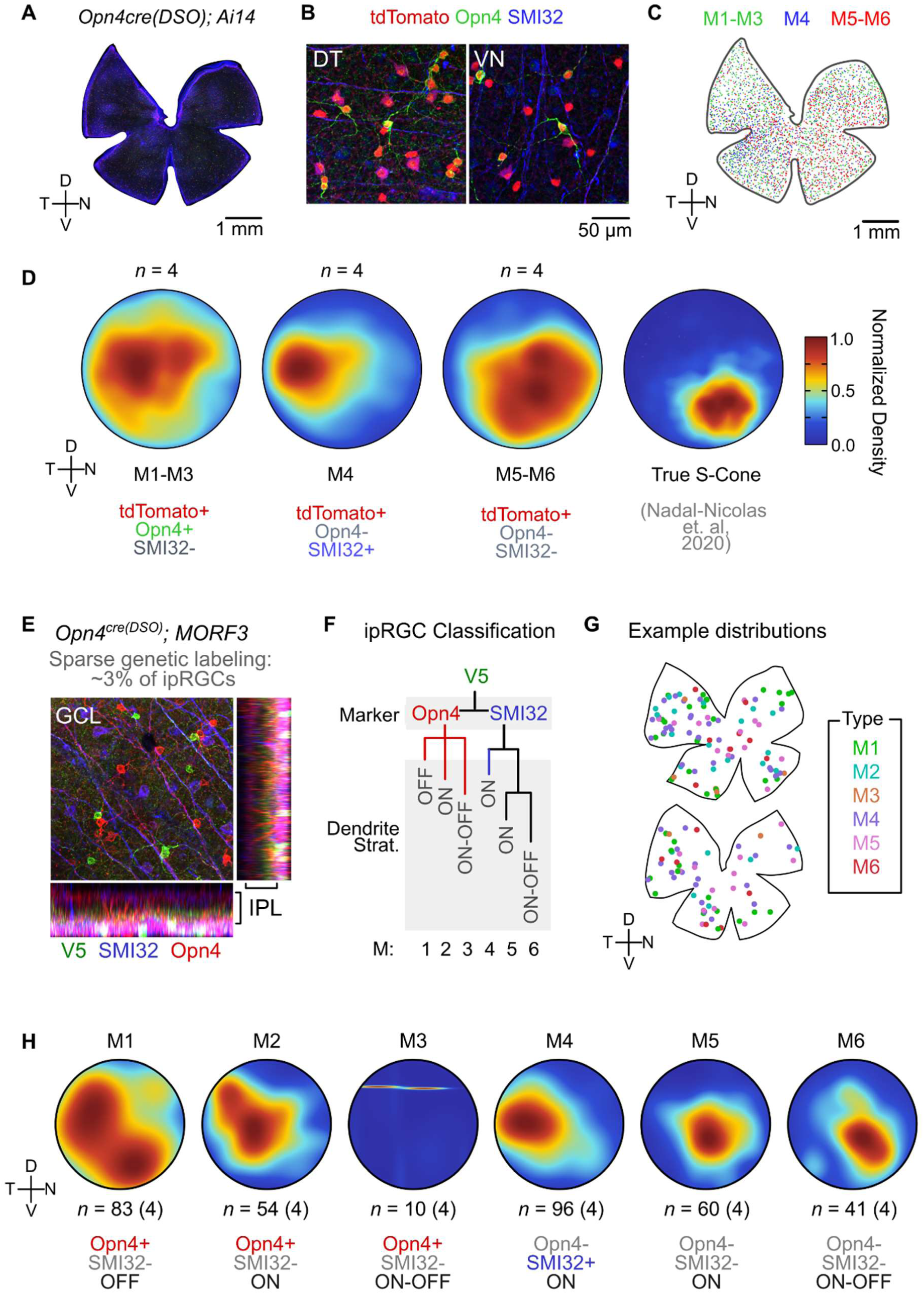
Spatial distribution of ipRGC types in the *Opn4cre(DSO*) mouse line. (A) Flat-mount representative image of *Opn4^cre(DSO)^; Ai14* stained against tdTomato (red), Opn4 (green), and SMI32 (blue). (B) Representative magnification of fields in the dorsotemporal (DT) and ventronasal (VN) locations highlighting labeling diversity. (C) Same flat-mount retina in (A) but with computed labels for each coarse ipRGC type – M1-M3 (tdTomato+ Opn4+ SMI32-), M4 (tdTomato+ Opn4-SMI32-), and M5-M6 (tdTomato+, Opn4-, SMI32-). (D) Normalized 2D density heatmaps from polar reconstructed retina, averaged across *n* = 4 per coarse ipRGC type, and analysis of data from Nadal-Nicolas et al., 2020 depicting true S-Cone distributions. (E) Representative high magnification confocal stack of genetic sparse labeling and multiple markers of ipRGCs, with profile projections. (F) Decision tree to define each ipRGC type using a combination of markers (Opn4 and SMI32) and dendritic stratification (ON, OFF, ON-OFF). (G) Representative flat-mount retinae with spatial locations of each ipRGC type used in this analysis. (H) Similar to (D) but for each ipRGC type (total number of cells *n* is located below each polar plot) averaged across number of retina shown in parentheses. GCL = ganglion cell layer, IPL = inner plexiform layer, D = dorsal, N = nasal, V = ventral, T = temporal.

Using this approach, we generated polar density maps (Fig 5d) that represent average spatial densities of each coarse ipRGC type from *n* = 4 animals (each polar plot can be found in S Fig 5a). Consistent with previous work, we found a broad distribution of M1-M3 ipRGCs, with a slight emphasis in the temporal retina^45^. Additionally, and similar to previous work on M4/Alpha ON-sustained RGCs, we find these cells in a strong nasal-to-temporal gradient, with highest density in the temporal retina^24,26^. Surprisingly, we find that M5 and M6 ipRGCs, while more broadly distributed than M4s, increase in density in the ventro-nasal (VN) retina (Fig 5d). We performed a similar analysis on published spatial coordinates for True S-Cones^47^ and find that the M5-M6 population is more spatially correlated to True S-Cones than to the M1-M3 or M4 types (S Fig 5b).

To more comprehensively define the spatial distributions of ipRGCs, we performed similar spatial analyses on sparsely labeled cells from the *Opn4^cre(DSO^*^)^; *MORF3* line. To categorize each V5-labeled ipRGC type, we used a combination of molecular markers (Fig 5e) coupled with dendritic stratification and morphology (Fig 5e-f). Thus, our criteria for ipRGC typing had 3 criteria: Opn4 expression (+/-), SMI32 labeling (+/-), and dendritic stratification (ON, OFF, ON-OFF) (see S Fig 5c for example types and reconstructions). In total, we surveyed 344 cells from *n =* 4 flattened retinae (Fig 5g) and analyzed their densities in polar space averaged across animals (Fig 5h). Consistent with our findings using *Ai14* and markers (Fig 5a-d), we find that:

- M1s are broadly distributed with a higher temporal density (83/344 cells).
- M2s are more restricted to the dorsal retina with a higher temporal density (56/344 cells)
- M3s are extremely rare (10/344 cells) and thus we could not conclude whether they are nonuniformly distributed
- M4s are strongly temporally enriched (96/344 cells), a finding identical to using tdTomato and SMI32 as markers in Fig 5a-d
- M5s and M6s appear to be enriched in the central-to-ventronasal retina (M5: 60/344, M6: 41/344)

Thus from these experiments we conclude that not only are all ipRGC types labeled in the *Opn4^cre(DSO)^* line, but that majority of types are nonuniformly distributed within the retina.

### ipRGC projections and *Opn4cre* labeling in the brain

ipRGCs collectively project to several central targets in the brain, the majority of which are involved in non-image forming vision (vLPO/POA, SCN, pSON, PHb, IGL, vLGN, IGL, OPN, PPT), with innervation of image forming nuclei as well (dLGN, SC)^48,49^ (Fig 6a). Previous work has shown that when crossed to *Ai14*, *Opn4^cre(Saha)^* labels several regions of the brain, including the targets of ipRGC innervation, the cortex, thalamic groups, and other nuclei^14,21^. While it is possible that *Opn4* could be expressed by several brain regions either developmentally or in the adult, given the already described off-target effects of this *Cre* line in the retina, we reasoned that further analysis of *Cre* labeling between lines was warranted.

**Figure 6.**
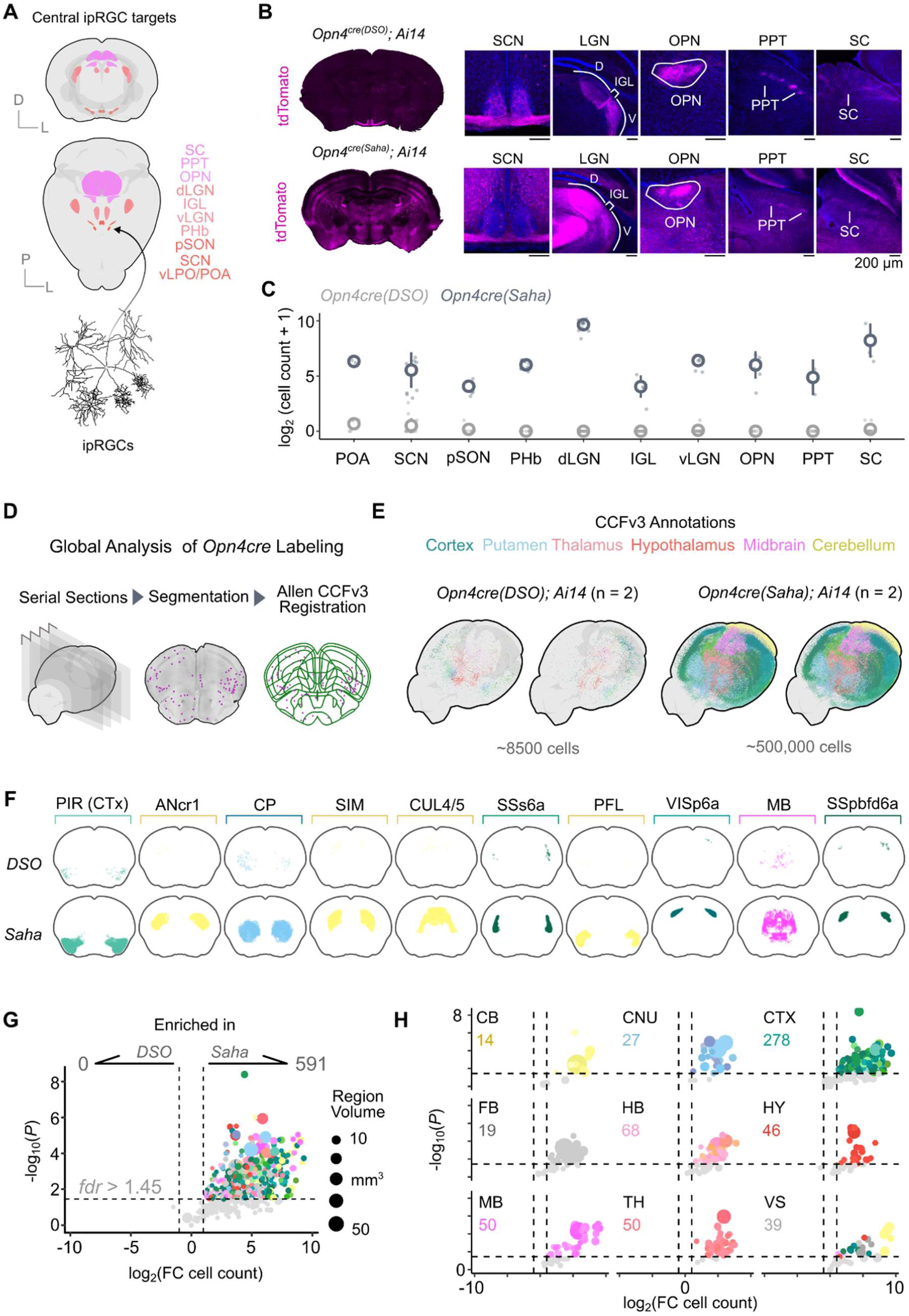
A brain-wide analysis of labeled cells reveals divergent labeling across Cre lines. (A) Volumetric projections of ipRGC targets in the Allen Brain Atlas CCFv3. (B) Representative sections and major ipRGC targets in the brain from *Opn4^cre(DSO)^; Ai14* (top) and *Opn4^cre(Saha)^; Ai14* (bottom) lines. (C) Quantification of absolute cell count, displayed as log_2_ (cell count + 1), for ipRGC targets between *Cre* lines. (D) Schematization of global cell labeling pipeline used in (E-H). (E) Whole brain point clouds of segmented cells in the *Opn4cre(DSO*) and *Opn4cre(Saha*) lines crossed to *Ai14*, colored by CCFv3 annotations. (F) Comparison between 10 regions with highest cell counts in the *Opn4cre(Saha*) line. (G) Volcano plot depicting differences in cell count per region with annotated cells. Numbers at the top depict number of regions in the CCFv3 with significant enrichment between the lines. (H) Same data in (G) but segregated by 9 parent structures. Abbreviations: POA = pre optic area, SCN = suprachiasmatic nucleus, pSON = peri-supraoptic nucleus, PHb = peri-habenula, dLGN = dorsal lateral geniculate nucleus, IGL = intergeniculate leaflet, vLGN = ventral lateral geniculate nucleus, OPN = olivary pretectal nucleus, PPT = posterior pretectal nucleus, SC = superior colliculus, PIR = piriform cortex, ANcr1 = crus 1, CP = caudoputamen, SIM = simple lobule, CUL4/5 = lobule IV-V, SSs6a = supplemental somatosensory area layer 6a, PFL = paraflocculus, VISp6a = primary visual area layer 6a, MB = midbrain, SSpbfd6a = Somatosensory barrel field associated area layer 6a, CB = cerebellum, CNU = caudoputamen, CTX = cortex, FB = fiber tracts, HB = hindbrain, HY = hypothalamus, MB = midbrain, TH = thalamus, VS = ventricle system

Using *Ai14* as a reporter, we assessed labeling in the primary targets of ipRGCs across both *Cre* lines. These analyses revealed strong tdTomato+ fibers in all central ipRGC targets within the *Opn4^cre(DSO^*^)^ line, consistent with the line’s specificity in the retina (Fig 6b top). Comparatively, the *Opn4^cre(Saha^*^)^ line suffered from major off-target cell body labeling in all targets assessed (Fig 6b bottom). While many cells were labeled in each central target within this line, the image-forming targets dLGN and SC had the highest number (Fig 6c). By contrast, in the *Opn4^cre(DSO^*^)^ line, while a few sporadic cells were detected unilaterally in the SCN (∼5-6 cells), no other cell bodies were located in ipRGC projection targets (Fig. 6b, c).

To establish a global map of labeling between these two lines, we generated serial sections of brains from 2.5 to -7.5 mm AP (anterior-posterior) across both lines crossed to *Ai14*, performed automated cell segmentation, and serially registered slices and cell coordinates to the Allen Mouse Brain Atlas using their Closed Coordinate Framework (CCFv3) (Fig 6d). Using this approach, we captured and registered the spatial locations of > 750,000 cells, and as such, only performed this analysis on *n =* 2 brains from each line (each mouse was from a separate litter). To validate that registration using this approach produced reproducible results, we split the dataset into odd and even serial section series and independently registered them to the CCFv3 (S Fig 6a), which showed high region-specific correlation, regardless of mouse line (S Fig 6b).

Using this approach, we identified cells in over 600 unique regions of the mouse brain, totaling ∼745,000 in the *Opn4^cre(Saha^*^)^ line. In comparison, 11,000 cells in total were detected in the *Opn4^cre(DSO^*^)^, representing ∼1.5% of the cells detected in the *Saha* line. In the *Opn4^cre(DSO^*^)^ line, cells appeared to be more enriched in the ventral brain, but appeared unilaterally, suggesting random recombination (Fig 6e, S Fig 7a). This is in contrast to the global distribution of cells in the *Opn4^cre(Saha^*^)^ line, which were found in all 7 parent brain structures within the CCFv3 (S Fig 7b). As a comparison, we computed the regions with the top 10 cell counts within the *Opn4^cre(Saha)^* line and a comparison between lines is presented in Fig 6f. Regions with the highest cell counts were within the cerebellum (4/10): Crus 1 (2^nd^), Simple Lobule (4^th^), Lobule IV-V (5^th^), and Paraflocculus (7^th^), followed by cortical structures (4/10): Piriform cortex (1^st^), Supplemental Somatosensory Area layer 6a (6^th^), Primary Visual Area layer 6a (8^th^) and Supplemental Somatosensory Barrel Field Associated layer 6a (10^th^). The caudoputamen (1/10, 3^rd^) and midbrain (1/10, 9^th^) were large structures that contained several thousand cells. In comparison, the *Opn4^cre(DSO^*^)^ line has minimal labeling in each of these regions, with bilateral asymmetries suggesting random recombination events (Fig 6f top row). Across 638 regions with cell bodies surveyed, 591 were significantly enriched in the *Opn4^cre(Saha^*^)^ line over the *Opn4^cre(DSO^*^)^ line when crossed to *Ai14* (Fig 6g). Almost half these regions represented cortical areas (CTX; 278/591), followed by hindbrain structures (HB; 68/591), midbrain (MB; 50/591), thalamus (TH; 50/591), hypothalamus (HY; 46/591), ventricular system (VS; 39/591), fiber tracts (FB; 19/591), and lastly cerebellum (CB; 14/591). No brain regions showed significant enrichment in the *Opn4^cre(DSO^*^)^ line when tested against the *Opn4^cre(Saha^*^)^ line. Assessment of cell bodies across the same volume from multiple litters of *Opn4^cre(DSO^*^)^; *Ai14* mice showed no consistent pattern of cell labeling, suggesting that *Opn4* expression is highly restricted to RGCs in the retina (data not shown).

**Figure 7.**
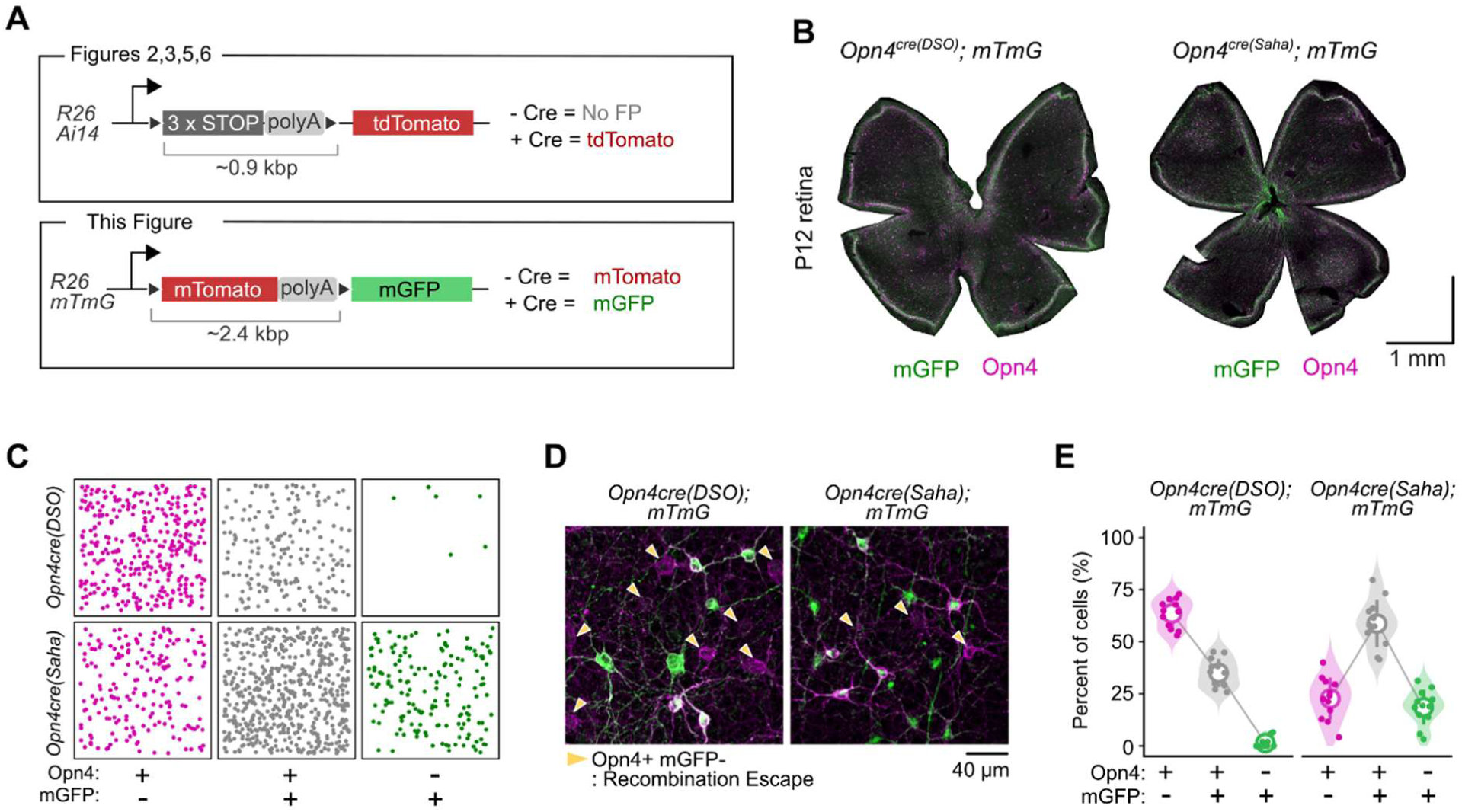
Reporter inter-*loxP* site distances augment recombination escape in *Opn4cre* lines. (A) Schematic depicting *Cre*-dependent reporters used in this study (top) and this figure (bottom). (B) Representative flat-mount retina from *Opn4cre(DSO)* and *Opn4cre(Saha*) crossed to *mTmG*. (C) Spatial point distributions of escaped cell (Opn4+ mGFP-; magenta), appropriately targeted cells (Opn4+ mGFP+; grey) and targeted cells (Opn4-mGFP+). (D) Representative images of analysis performed in (C & E). Arrowheads depict escaped cells (Opn4+ mGFP-). Mixed plots depicting percent of all analyzed cells in each category, across both lines. Small points = individual FOVs, large circles = mean of group, error bars = standard deviation.

### Altering inter-*loxP* distances shifts recombination efficiency of *Opn4^cre^* lines

The recombination efficiency of *Cre* lines depends on several factors including *Cre* expression level, genomic accessibility at the *loxP*-flanked locus, and the inter-*loxP* distance^23,50,51^. The *Ai9* and *Ai14* series of reporters are sensitive because of their CAG promoter-driven expression of tdTomato is within the highly accessible *ROSA26* locus^20^. With this genomic context, we can observe stark differences in recombination between both *Opn4^cre^* lines in both the retina and brain.

When *Opn4^cre(Saha^*^)^ is crossed to *Z/EG* or *Z/AP,* as done for most analysis using this line, labeling is more specific within the retina than compared with *Ai9* or *Ai14,* but the sensitivity of *Cre* labeling is diminished (∼30% Opn4+ cells escape recombination)^21,22^. Even with this enhanced specificity, alternative cell types remain labeled in the retina (rods and cones) and in the brain (dLGN, IGL, vLGN, Piriform cortex; See S Fig 4 in Ecker et al., 2010)^14^. These differences in labeling likely occur for 2 major reasons. The first is that the *Z/EG* and *Z/AP* alleles are both within regions of unknown accessibility in the genome^52,53^. Both lines show differences in the cell types that they label under ubiquitous *Cre* drivers, suggesting the local genomic architecture can modify expression of their reporters^52^. The second is that both transgenic lines require recombination of a cassette with large inter-*loxP* distances (>4.5kb), containing a lacZ-pgkNeoR-polyA sequence between *loxP* sites^52^. The combination of these variables likely means strongly reduced efficiency of even a boosted *Cre* line.

We set out to determine how altering components of a *Cre*-dependent reporter system would influence recombination efficiencies in both *Opn4 Cre* lines. To this end, we utilized the *mTmG* line that contains a *loxP-mTomato-polyA-loxP* (∼2.4kb) upstream of a mGFP coding sequence (Fig 7A). While this has a larger inter-*loxP* distance than *Ai14* (∼0.9kb), it has identical accessibility as it is in the *ROSA26* locus (Fig 7a). We crossed *Opn4^cre(DSO)^* and *Opn4^cre(Saha^*^)^ mice to the *mTmG* line, labeled retina for mGFP and Opn4 as in the *Ai14* experiments (Fig 2) and assessed recombination efficiency (Fig 7b-e). We performed these experiments on P12 mice, as M4s express higher levels of melanopsin during development (Fig 7b)^54^. We characterized all Opn4+, mGFP+ cells, and their overlap, categorizing them into (1) cells that escaped recombination (Opn4+ mGFP-), (2) cells that were accurately targeted (Opn4+ mGFP+), and non-Opn4 expressing cells (Opn4-mGFP+) which could be ipRGCs and or off-target labeling (Fig 7c,d).

We found that, in contrast to our experiments with *Ai14*, the *Opn4^cre(DSO)^* line displayed significant recombination escape (Opn4+ mGFP-; 63.6 ± 6.8%), while the remaining cells were appropriately targeted (Opn4+ mGFP+; 35.0 ± 6.8%), and almost no cells contained mGFP only (Opn4-mGFP+; 1.36 ± 2.4%; Fig 7e). In contrast, in the *Opn4^cre(Saha^*^)^ line, only 22.7 ± 10.1 % of cells escaped recombination (Opn4+ mGFP-), while 58.7 ± 11.4 % expressed both Opn4 and mGFP, and the remaining 18.6 ± 8.3 % of cells expressed mGFP only (Fig 7e). Thus, both *Opn4^cre^* lines are affected by increasing inter-*loxP* site distances of floxed constructs, even at a highly accessible locus.

## DISCUSSION

In this study, we generated a new *Opn4^cre^* knock-in mouse line (*Opn4^cre(DSO)^*) and provide an in-depth comparison between it and the most widely used ipRGC-targeting *Cre* line (*Opn4^cre(Saha)^*). By placing *Cre* immediately downstream of the *Opn4* start codon and using the native *Opn4* promoter to drive recombinase expression, we found that ipRGC photoreceptors within the retina can be efficiently and selectively labeled when crossed to widely used, chromatin accessible *Cre-*dependent reporters (*Ai9*, *Ai14*, and *MORF3;* Fig 2-6). Owing to the enhanced specificity of the *Opn4^cre(DSO^*^)^ line, we uncovered the spatial distributions of M1-M6 ipRGCs using both molecular markers and sparse labeling, and found that the M2, M4, M5, and M6 types are nonuniformly distributed across the retina. In contrast, we confirmed prior reports that the *Opn4^cre(Saha)^* line labels ipRGCs, but these cells represent a small minority of the total labeled cells across the central nervous system. It is likely that the distinct design of these two alleles leads to vastly different labeling within the eye and brain.

### Retinal *Cre* efficiency and off-target cells

Our assessment of both *Cre* lines in the retina revealed two important findings. The first is that both *Cre* lines, even with differing regulatory elements, are similarly efficient at labeling ipRGCs expressing detectable melanopsin protein (∼99%, Fig 2). This is surprising given that widespread recombination in non-ipRGC types occurs in the *Opn4^cre(Saha^*^)^ line, suggesting a negligible gain in sensitivity by boosting *Cre* expression coupled to sensitive reporters. Secondly, while it has been noted that the *Opn4^cre(Saha^*^)^ line labels non-ipRGCs in the outer retina (S Fig 1 & S Fig 2), it is apparent that it labels other types of RGCs and some amacrine cells in the inner retina as well (S Fig 2). This would explain why most cells lack an intrinsic photocurrent in *Opn4^cre(Saha)^; Ai14* mice^21^. Importantly, this does not appear to be restricted to crosses with sensitive reporters. The original paper detailing the *Opn4^cre(Saha^*^)^ line reported that ∼10% of recorded cells lacked an intrinsic photocurrent. This may be attributable to recombination of *Z/EG* in non-ipRGCs or non-RGCs within the inner retina^14^.

Given the variable expression of melanopsin between ipRGC types, it is difficult to estimate the true sensitivity of either *Cre* line. For this, we must be able to independently label each type of cell and describe the overlap between them and *Cre*-dependent reporters. Based on previously reported counts of Opn4+ cells from various reports, both lines appear to be sensitive enough to label the entire population (Fig 2a-f)^25,27–30^. However, it is unclear how sensitive they are to label or recombine *loxP*-flanked cassettes in M4-M6s. Surveying the entire retina for M4s using the *Opn4^cre(DSO^*^)^; *Ai14* (tdTomato+ SMI32+, Fig 5a-c), we found 722 ± 58 cells, representing ∼1.44% of all RGCs similar to the reported ∼840 Alpha ON-sustained cells per retina^55^. However, with no alternative means to label the M5 and M6 population comprehensively and independently of *Opn4^cre(DSO^*^)^, it remains unclear if this, or any line captures the majority of these low melanopsin-expressing populations.

### Central-to-ventronasal enrichment of M5 and M6 types

Even without metrics of absolute sensitivity, the utility of this new *Cre* line is clear: even with low expression of melanopsin, it is capable of labeling M5s and M6s as a group (Fig 5a-c), or individually and stochastically in sufficient quantities for spatial analysis (Fig 5g). We used this feature to study the nonuniform spatial distribution of each ipRGC type to find that both the M5 and M6 type are spatially enriched in the ventronasal retina. This pattern is reminiscent of developmental *Cdh3-GFP* labeling in the retina, a marker known to label both M5 and M6 ipRGCs^40,46^. We also show empirically that their spatial density patterns strongly correlate with the true S-Cones^47^ that are enriched in the ventronasal retina and likely serve as the outer sensory basis of M5 color opponency (UV-ON, green-OFF)^39^. Similarly, type 9 bipolar cells (or S-Cone Bipolar Cells; SCBCs) which receive input from true S-Cones and provide input to M5s are enriched in a similar spatial gradient^39,47,56^. Together, these data highlight several components of a microcircuit displaying spatial anisotropies within the retina.

In contrast, the M6 type shares spatial enrichment with both true S-Cones and M5s, but does not appear to encode chromatic information^40^. This spatial enrichment is observed in lines such as the *Cdh3-GFP* and in experiments tracing RGCs projecting to the olivary pretectal nucleus (OPN), which is dominated by M6s^57^. One potential explanation for this consistent spatial pattern, but lack of color opponency may be the differences in bipolar inputs to M6 (Type 2 and 9o) compared to M5^58^. Further characterization of the microcircuits and optimal responses of M6s will be necessary to understand why they are localized to the ventronasal retina but encode different visual features than their M5 counterparts.

### A case for selective expression of *Opn4* in the central nervous system

Photopigments are expressed in cells beyond the neural retina and the eye^59^. For example, *Opn3* and *Opn5* are distributed in tissues outside the eye and have had reported functions within the brain^60–62^. Expression of *Opn4* across the brain would be consistent with other photopigment expression patterns and potential function. However, given that the *Opn4^cre(Saha^*^)^ line labels many regions, including those that are targets of the ipRGCs themselves, it is hard to accurately determine whether any bona fide expression of *Opn4* takes place elsewhere in the central nervous system.

Using the *Opn4^cre(DSO)^* line, we found limited evidence for expression within the brain. While we see sporadic, often unilateral recombination of cells in different locations, this is never consistent. This likely represent a feature of the sensitivity of *Ai14* as a reporter, rather than an underlying expression pattern of *Opn4*. Similar features have been seen with other photopigment *Cre* lines in the brain, with uni-hemispheric labeling in regions unrelated to normal expression^60^. While our whole brain analyses focused on brain volumes between 2.5 mm to 7.5 mm (AP), and could have limited sampling beyond these bounds, we never encountered fiber tracts or terminals that terminate within the volumes assessed. This suggests that at least no long-range projecting neuron types express *Opn4*. We did not, however, survey the rest of the organism for this study. Other groups have reported functions of melanopsin in the trigeminal nerve^63^, iris sphincter muscle^64^, vascular relaxation^65^, and adipocyte physiology^66^, and further work will be necessary to determine whether these tissues are labeled by the *Opn4^cre(DSO)^* line.

### Implications of *Cre* use for conditional knockout experiments

While we performed an in-depth analysis of labeling using various reporter lines, the conclusions of these experiments have implications that reach beyond tagging cells for visualization. The *Opn4^cre(Saha^*^)^ line likely drives strong *Cre* expression but at the apparent expense of specificity. This is observed across sensitive reporters such as *Ai9* and *Ai14*, moderately sensitive reporters such as *mTmG*, and fairly insensitive reporters like *Z/EG*. Importantly, this can have an impact on experiments assessing function and behavior that utilize *Opn4cre(Saha*) as a driver to abolish gene expression conditionally (*Slc17a6^flox^*, *Gad1/2^flox^*)^18,67^ or drive suicide gene expression (*Pou4f2^zDTA^*)^17^. Given that both genomic accessibility and inter-*loxP* distances contribute to *Cre* recombination efficiency^23,68,69^, experimental work using this line must be wary of deleterious off-target recombination of floxed cassettes across the eye and brain.

We argue that experiments utilizing the *Opn4^cre(Saha^*^)^ line to abolish gene expression must assess not only the efficiency of deletion in the retina with methods such as RNA-scope and immunofluorescence, but also characterize recombination across the eye and brain. As typical reporters and floxed alleles of genes are differentially accessible and vary in their inter-*loxP* distances, a reporter cannot substitute for determining if gene expression has been specifically abolished and if off-target events have occurred^70^. While it is possible that the current *Cre*-dependent reporters oversample due to their ease of genomic access and small inter-*loxP* distances, certain widely used lines such as *Slc17a6* (encoding Vglut2) and *Gad2* (GAD65) have flox lines with short inter-*loxP* distances – Floxed Exon 1 of *Scl17a6^flox^* is < 0.5kb^71^ and floxed Exon 1 of *Gad2^flox^* is < 0.1kb^72^. Both genes are highly accessible in excitatory and inhibitory neurons, due to the very nature of the gene product, and as such will be susceptible to off-target recombination. Given that over 591 regions of the brain were labeled in the *Opn4^cre(Saha^*^)^ line, investigators must consider potential confounding effects in their experiments, especially at the organismal and behavioral level.

Conversely, while our line is more specific with *Cre*-dependent reporters, because of the variable and low expression of the *Opn4* locus, recombination efficiency may be limited for genes with large inter-*loxP* site distances or in regions of the genome with lower accessibility. Thus, for conditional loss-of-function experiments, we recommend assessing ipRGCs for successful recombination using *in situ* transcript detection or immunohistological approaches to verify that generalized loss-of-function has taken place. Moving forward as a field, we must continue to explore methods to genetically access ipRGC types using gene drivers that are both highly expressed and cell type specific, which would allow us to circumvent using the variably expressed *Opn4* locus to interrogate ipRGC function in the eye and in behavior.

## CONCLUSION

We developed a new *Opn4^cre^* recombinase that is driven by native regulatory elements of the *Opn4* locus. While the expression of melanopsin (Opn4) varies dramatically between ipRGC types within the retina, when crossed to *Ai9, Ai14,* and *MORF3, Opn4^cre(DSO)^* is capable of selectively labeling all ipRGC types with minimal off-target labeling. By providing an in-depth comparison between recombination between it and a widely used existing *Opn4^cre(Saha^*^)^ line, we highlight the need to characterize recombination efficiencies and off-target effects prior to performing experiments.

## MATERIALS & METHODS

### Mice

#### Mouse line generation

The *Opn4^cre(DSO)^* mouse line was generated in-house at the Cincinnati Children’s Hospital Transgenic Core using CRISPR-Cas9 targeting and long single strand DNA (ssDNA) knock-in technology. A single guide RNA (gRNA) targeting the region between the 5’ UTR and Exon 1 of the mouse *Opn4* locus (mm10; cut site Chr14:34,599,959) was generated and validated in ES cells for cutting efficiency. A long ssDNA donor containing 5’ and 3’ homology arms (∼370 bp), followed by *Cre* recombinase, and SV40 polyA signal was synthesized to generate *Opn4^cre(DSO)^* knock-in mice. Pronuclear injections of gRNA, long ssDNA donor, and Cas9 were performed on fertilized C57BL/6J (JAX #000664) eggs. From this, 1 male was generated with the appropriate insertion which was confirmed via insertion-specific PCR validation and subsequently targeted sequencing of the insert.

gRNA sequence (**PAM**): 5’-CTGAAGGAGAGTCCATGCTC **AGG-**3’

#### Mouse lines

Mice between the ages of P30-P60 (both sexes) were used in this study, except for P12 mice in Fig 7 which is specified in text and figure. All animals were housed and maintained in a pathogen-free vivarium in accordance with protocols approved by the Institutional Animal Care and Use Committee (IACUC) at Cincinnati Children’s Hospital Medical Center and Washington State University. Other genetically modified mouse lines used in this include: *Opn4^cre(Saha)^* (*Opn4^tm1(cre)Saha^/J*; #035925), *Ai9* (*Gt(ROSA)26Sor^tm9(CAG-tdTomato)Hze^*/J; #007909), *Ai14* (*Gt(ROSA)26Sor^tm^*^14^*^(CAG-tdTomato)Hze^*/J; #007914), *MORF3* (*Gt(ROSA)26Sor^tm3(CAG-sfGFP*)Xwy^*/J; #035403), and *mTmG* (*Gt(ROSA)26Sor^tm4(ACTB-Tomato,-EGFP)Luo^/*YgchJ; #037456).

### Histology and Tissue Processing

#### Tissue Processing

Retinae and brains were processed as previously described. Briefly, whole eyes were fixed in 4% paraformaldehyde (PFA) for 40 minutes at room temperature, followed by a PBS wash. Retinae were dissected out and cryoprotected (for sectioning) or processed for whole mount immunofluorescence. Brains were processed similarly, with fixation in 4% PFA overnight at 4C, followed by 2x washes with PBS and serial dehydration/cryoprotection. Brains were serially sectioned at 40 µm thickness, adhered directly to SuperFrost charged slides, and stored at -20C until analysis.

#### Histology and Immunofluorescence

Retina and brain were permeabilized in 0.5% Triton-X 100 in PBS (PBST), followed by blocking in 10% normal donkey serum diluted in PBST. Sections were incubated in primary antibodies (see table below) overnight, with the exception of wholemount retina, which were stained on an orbital rocker at 4°C for 2-3 days. Following primary incubation, tissue sections and wholemounts were washed up to 6 times with PBS, followed by re-permeabilization with 0.5% PBST, and a 1-hour incubation with secondary antibodies (1:1000 dilution) at room temperature. Following 6+ washes with PBS, tissue sections and whole retina were cover-slipped and imaged the same day.

#### Primary and Secondary antibodies

The following primary antibodies were used in this study: anti-tdTomato (SICGEN, #AB8181, RRID:AB_2722750), anti-Melanopsin N15 (a gift from Ignacio Provencio), anti-Melanopsin (Advanced Targeting Systems, #AB-N39, RRID:AB_1608076), anti-Rbpms (abcam, #ab152101, RRID:AB_2923082), anti-VAChT (Synaptic Systems, #139103, RRID:AB_887864), anti-Tbr2/Eomes (abcam, #ab183991, RRID:AB_2721040), anti-Calb2 (abcam, #ab277631, RRID: N/A), anti-V5 (Novus Biological, #NB600-379, RRID: AB_10003214), SMI32 (Enzo LifeScience, #ENZ-ABS219, RRID: N/A), anti-GFP (Novus, #NB100-1614, RRID: AB_10001164). The following secondary antibodies were used (Jackson ImmunoResearch): Donkey anti-rabbit 488 (#711-545-152) & 647 (#711-605-152), Donkey anti-chicken 488 (#703-545-155), Donkey anti-mouse 647 (#715-605-151), and Donkey anti-sheep 594 (#713-585-147).

### Cell counting and spatial analyses

When possible, automated, and validated methods were employed to count and annotate cell positions on images (cellpose, see below). However, certain experiments with low signal-to-noise or multiple markers that did not mark the same cellular domain (tdTomato, SMI32, melanopsin) required manual annotation as downstream analytical methods could not reliably distinguish different populations computationally.

#### Manual cell counting and annotation

For experiments in Fig 3 & Fig 5, cells were manually annotated by importing confocal images into FIJI, followed by annotation using the CellCounter plugin. Subsequently, an XML file corresponding to the spatial coordinates and a csv file corresponding to the features (x, y, Area, etc.) of the annotated cell were exported and used from downstream analysis or further processing (density, etc.).

#### Automated cell segmentation

To segment cell boundaries for downstream analysis, we utilized cellpose, a convolutional neural network trained on a variety of cellular bioimage data. For tdTomato cell segmentation, we utilized a pretrained model (cyto) with parameters cell_diameter = 12.5, cellprob_threshold = -1.0 and flow_threshold = 0.4. For Opn4, we used an alternative pretrained model (cyto2) with default parameters. Each image was inspected following automated segmentation for errors and dropouts, subsequently corrected in the built-in GUI, and exported as a text file with polygon coordinates for each cell.

#### Cell type classification following automated segmentation

To determine if cells overlapped in marker expression, as performed in Fig 2, we used a spatial nearest neighbor approach with stringent criteria. First we imported polygons as (x,y) coordinates for each marker (tdTomato, Melanopsin) per image. Then we computed the centroid position of each cell in 2D space using their polygons. To determine if masks overlapped and thus represent double positive cells (tdTomato+ Opn4+), we computed the nearest neighbor distances between different markers and assigned masks to be shared between cells if their centroid distances were less than 4 pixels (nnd_tdTomato_ ◊nnd_Opn4_ < 4). Using this method, we estimated the total number of tdTomato+ Opn4+ cells, tdTomato+ Opn4-cells, and tdTomato-Opn4+ cells for each retina analyzed. We confirmed that this method accurately captures overlapping and non-overlapping masks by employing two different quality control measures: (1) for a random sampling of retina we manually annotated after running this analysis pipeline, and independently assessed cell type assignments projected onto each retina and found nearly identical results. (2) Following automated type assignment, we visually inspected each sample with its cognate image to be sure that masks were not spuriously assigned to each group.

#### Single cell transcriptome mining

Single cell transcriptomes of atlas RGCs (Tran et al., 2019) were downloaded from the Gene Expression Omnibus (GEO) under GSE137400. We performed standard preprocessing by including only genes expressed in >3 cells and cells with >200 expressed genes. Following this general preprocessing, we next performed integration using the Seurat v4 pipeline to eliminate batch effects. For this, we split the experiment by their sample id (aRGC1-9) and batch (1-3), scaled and transformed the data using the SCTransform() function while regressing mitochondrial genes. Next, we generated anchor features using SelectIntegrationFeatures() using 3000 variable features. The data was then integrated, using the default parameters, and preprocessed using a standard Seurat v4 workflow. Next, we subset the ipRGCs (and related clusters) using corresponding metadata from the original paper and performed dimensionality reduction and UMAP embedding (for visualization) and differential gene expression analysis to define markers of ipRGC types.

### Electrophysiology

Methods for path-clamp recordings from ipRGCs were similar to methods previously described_73_. Following at least one hour of dark adaptation, mice were euthanized by cervical dislocation after isoflurane anesthesia. Eyes were enucleated under dim red light, and the cornea was slit to remove the lens, followed by retina extraction into oxygenated, bicarbonate-buffered Ames media (US Biologicals), 3 mM kynurenic acid (Sigma), 50 µM DL-AP4 (Tocris), and 50 µM (+)-bicuculline (Sigma). The retinas were then cut into quarters, mounted on a nitrocellulose filter with a hole providing access to the retinal ganglion cell layer, which were stored in the same solution in the dark at room temperature. Retinas could be maintained for more than 4 hours under these conditions before recording.

#### Patch-Clamp Recordings

Whole-mounted retinas on nitrocellulose were transferred to the recording chamber, where they were maintained in bubbled Ames’ media (35C) flowing at 2.5 mL min^-1^. Retinal ganglion cells were visualized with infrared Dodt gradient contrast optics through an inverted Olympus microscope (BX51WI) using a 40X water-immersion objective, a Nikon DS-QiMc camera, and NIS Elements Advanced Research Imaging software (v3.20; Nikon). The microscope was mounted on a Sutter X-Y translation stage; electrode positioning was accomplished via a micromanipulator (MP-285; Sutter). Micropipettes were fabricated from filamented borosilicate glass with resistances of 5-6 MOhm when filled with a potassium-based internal solution containing (in mM) 125 K-gluconate, 2 CaCl_2_, 2 MgCl_2_, 10 EGTA, 10 K-HEPES, 0.5 Na_2_-GTP, 2 Mg-ATP and 0.3% Neurobiotin (pH 7.2). Synaptic blockers, which allowed for the isolation of intrinsic melanopsin-based photocurrents, were added to Ames’ media to the following concentrations: 3 mM kynurenic acid (Sigma), 50 µM DL-AP4 (Tocris), and 50 µM (+)-bicuculline (Sigma; from a 1000X stock in DMSO). ipRGCs were identified by tdTomato fluorescence using a short exposure to 540 nm light (<1s; 1 x 10^6^ photons cm^-2^ s^-1^; Lambda DG-4, Sutter Instruments).

For light response experiments, recordings were made in voltage-clamp mode at a holding potential of -60 mV (not corrected for a junction potential of -13 mV). Responses were low-pass filtered at 5Hz post hoc (SigmaPlot) to reduce noise and breakthrough action potentials. Maximal melanopsin-based intrinsic photocurrents were elicited with a 100 ms 480 nm blue-light stimulus delivered from a Lambda DG-4 light source (Sutter Instruments) with an intensity of 2 x 10^10^ photons um^-2^ s^-1^. The intensity of background illumination from the computer monitors throughout these experiments was no greater than 1 x 10^-9^ W cm^-2^.

#### Immunohistochemistry post recording

After patch-clamp recording from an ipRGC, the retina was fixed in 5% paraformaldehyde for 1 hour and washed 3 times with 1 x PBS for 20 minutes before applying immunohistochemistry buffer (0.5% Triton X-1000, 1% BSA, 5% Normal Donkey Serum in PBS). Immunohistochemistry was performed using rabbit anti-Melanopsin (mOpn4)^74^ and streptavidin-488 (1:1000, Invitrogen). After immunostaining, the retinas were mounted and imaged using a Leica White Light Laser Confocal Microscope (TCS SP8 X).

#### Single ipRGC reconstructions

High resolution images of cells labeled in the *Opn4^cre(DSO)^; Morf3* retina were acquired using a 20x objective and 2x digital zoom (z-step 0.2um) and traced semi-automatically using the Neuroanatomy toolbox (formerly Simple Neurite Tracer; SNT) in ImageJ. Briefly, single channel z-stacks were imported into SNT, a root was set at the center of the soma and using the NBA*Search tracing algorithm, positions were defined along the cell that branched from the soma root. Cells were continually assessed in both the XZ and YZ planes to ensure no artifacts were introduced during reconstruction. Reconstructed cells were saved as SWC files to plot in XY and XZ dimensions using the natverse library (v1.8.19) in R.

### Spatial and Polar reconstruction of whole mount retina

#### Polar reconstruction

To project cells from wholemount images to polar, retinotopic space, we implemented the retistruct package in R (v0.6.3). For these experiments, a deep dorsal cut was made during dissection to identify the dorsal pole and left/right retina were noted and separated for analysis. (*x,y*) coordinates of cells across the entire retina and the retinal outline were acquired as per Sterratt et al., and retina were reconstructed using *φ_0_* = 115° (based on adult mouse ocular properties in Sterratt et al.). The newly reconstructed retina had each cell position (*x,y*) converted into *phi* and *lambda* polar coordinates, which were subsequently saved and used for downstream analysis.

#### Spatial density analysis

By performing the previous step, we can register cellular coordinates to common reference space and leveraged this to understand the spatial heterogeneities of different ipRGC types. To study the spatial densities of cells, we performed kernel density estimation (KDE) on the 2D polar coordinates using the MASS package in R. For this, we first estimated the joint range of cell type *phi* and *lambda* positions, then performed 2D-KDE using the kde2d() function with n=100 grid points in each direction. The output of this analysis is probability densities for each cell type across a reference retinal grid (as in Fig 5).

### Whole brain lineage tracing analysis

#### Automated cell segmentation

Similar to analysis performed in the retina, to automatically segment cells from serial sections of brain from *Opn4^cre(DSO)^; Ai14* and *Opn4^cre(Saha)^; Ai14* mice, we utilized cellpose’s cyto model with parameters flow_threshold = 0.4, cellprob_threshold = -1, and cell_diameter = 12.5. Images were batch processed using scripting and directories of text files with polygons corresponding to individual cells’ masks were used for downstream analysis. These masks were then converted to centroid *(x,y)* coordinates in image-space that required registration to the Allen Mouse Brain Atlas Common Coordinate Framework v3 (CCFv3).

#### Manual registration of sections

To register sections and their cognate cell positions to the CCFv3, we utilized a MATLAB pipeline developed by the Root lab^75^. Briefly, this pipeline allows for individual sections to be registered to the 10 µm atlas in CCFv3 with flexibility in the cutting angle of the section, as the atlas volume can be sliced in 3-dimensions (AP, DV, ML). Using this method, we independently registered odd and even series of brain sections from both mouse lines to validate that this approach yields consistent and reproducible results. After slices and cell coordinates had been transformed, cells are assigned a region-specific label based on their location in atlas-space and were registered spatially to the CCFv3.

### Analysis, Statistics, and Reproducibility

#### Investigator blinding

Investigators were not blinded to the genotypes and sexes of the animals used in this study. However, most analyses comparing cell counts between genetic lines utilized automated approaches that meant limited input from investigators.

#### IPL lamination of dendrites

To investigate the stratification of *Opn4^cre^*-labeled cells in the IPL, we imaged the inner plexiform layer (IPL) at 40x resolution, generated ∼5 55x35 µm ROIs centered within the IPL and aligned with the IPL-INL boundary. We then extracted mean intensity values for each channel (tdTomato, VAChT) across the IPL depth (55 µm). Each ROI channel was normalized to their maximum intensity value for downstream analysis and comparison. The analysis included 49 *Opn4^cre(DSO)^* ROIs (*n* = 4 animals) and 46 *Opn4^cre(Saha)^*ROIs (*n* = 4 animals).

#### Clustering ipRGC photocurrents

To cluster intrinsic photocurrents produced by cells during whole-cell recording, we limited our analysis to time points corresponding to -1 and 30 seconds from stimulus onset. Additionally, we down-sampled recordings to 50Hz for computational efficiency. Currents were converted to a matrix that was centered to the cell’s mean response and scaled to unit variance prior to clustering. Then, using the hclust() function with nstart = 50, we performed hierarchical clustering on photocurrents. To determine optimal clusters, we used the gap statistics method in the factoextra package (v1.0.7).

#### Average ipRGC spatial densities and correlation

Following KDE analysis on individual ipRGC types from each retinal sample, given that KDE produces a grid with probability densities, we leveraged this to (1) generate an average map of broad ipRGC type densities (Fig 5) and (2) use this density-coordinate grid to correlate different types of ipRGCs and other photoreceptors (True S-Cones). To generate average density maps, first each sample was scaled by subtracting their minimum spatial density followed by dividing by their new maximum density, transforming the density bounds to (0,1). This was repeated across each cell type per sample and used for subsequent analysis. Similar analyses were performed on sparsely labeled ipRGCs from the *Opn4^cre(DSO)^; MORF3* line. True S-Cone spatial distributions (x,y coordinates) were downloaded from Nadal-Nicolás et al., where a variant of polar reconstruction had been performed, and processed through our analysis pipeline. To correlate spatial densities of different cell types, we performed Spearman’s rank correlation between each of the normalized densities per sample. The results from this analysis are visualized as a clustered heatmap in Supplemental Fig. 4.

#### Statistical Analysis

All statistical analyses were performed in R using the rstatix package (v0.7.0). Statistical tests and corresponding sample sizes (*n*) are noted in the figure legends. For all analyses utilizing multiple comparisons, or multiple independent tests of a hypothesis, *p-*values were corrected via false discovery rate correction (*fdr* in text). Sample sizes were not predetermined.

## Resource Availability and Author Contributions

Further information and requests for resources and reagents should be directed to and will be fulfilled by the lead contact, Shane D’Souza

### Materials Availability

The *Opn4cre(DSO*) mouse line is available upon request from the lead contact. This mouse line will also be deposited to JAX, interest permitting.

### Data and Code Availability

All data and code used in this paper will be shared by the lead contact upon request after publication.

## Acknowledgements

We thank Paul Speeg for his excellent colony management. This work was supported by 1R01EY034456 to RAL, 1R01EY029202 to RLB, funds from the Abrahamson Pediatric Eye Institute (CCHMC) to RAL, the WSU-CVM Marvel Shields Autzen Fund to RLB, and the Albert J. Ryan Fellowship to SPD.

## Author Contributions

SPD conceived and directed the study. BD and SPD performed experiments and computational analysis. SOY and RLB performed patch-clamp and dye-filling experiments. RAL and SPD provided project leadership. SPD wrote the manuscript. BD, SOY, RLB, RAL, and SPD edited the manuscript.

## SUPPLEMENTAL FIGURES & LEGENDS

**Supplemental Figure 1.**
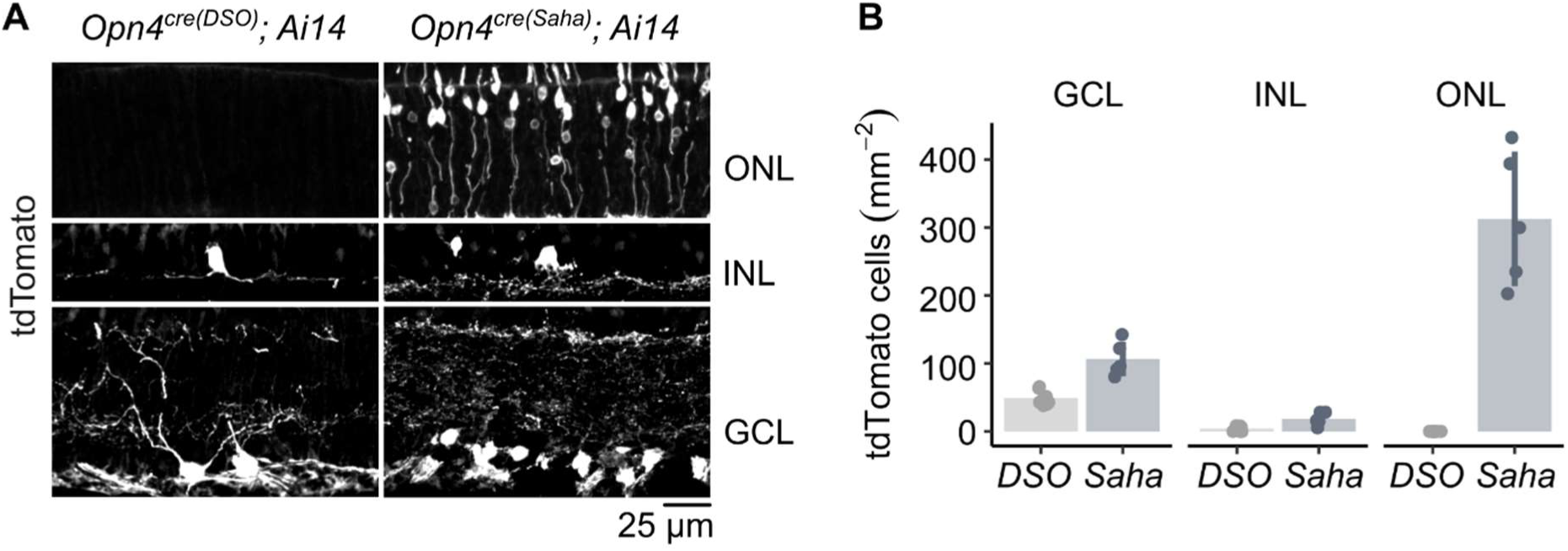
Labeled cell densities across retinal nuclear layers. (A) Representative confocal images highlighting tdTomato+ cells within the GCL, INL, and ONL between two *Cre* lines. (B) Quantification of densities (tdTomato+ cells / area) of images like in (A). Bar height = means, error bars = standard deviations. GCL = ganglion cell layer, INL = inner nuclear layer, ONL = outer nuclear layer.

**Supplemental Figure 2.**
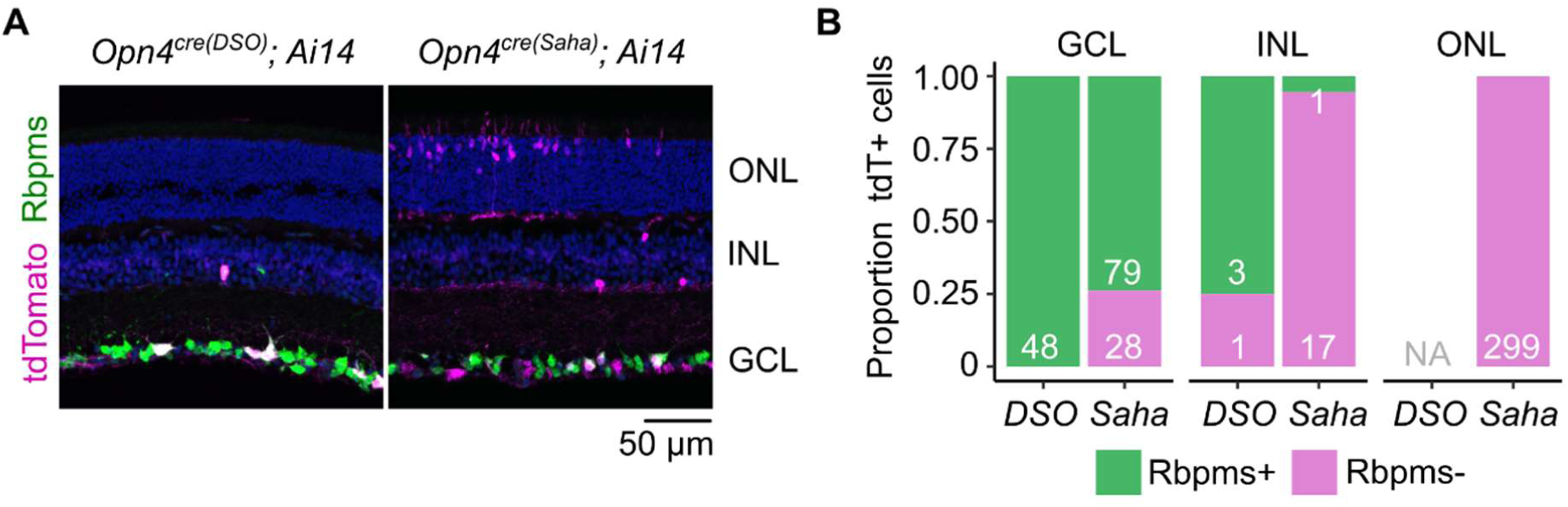
Retinal ganglion cell specificity between both lines. (A) Representative confocal images highlighting tdTomato+ and Rbpms+ cells within the GCL, INL, and ONL between two *Cre* lines. (B) Stacked bar graphs representing percentage of tdTomato cells that are Rbpms+ (green) and Rbpms-(magenta), numbers in each bar indicate number of cells surveyed. GCL = ganglion cell layer, INL = inner nuclear layer, ONL = outer nuclear layer.

**Supplemental Figure 3.**
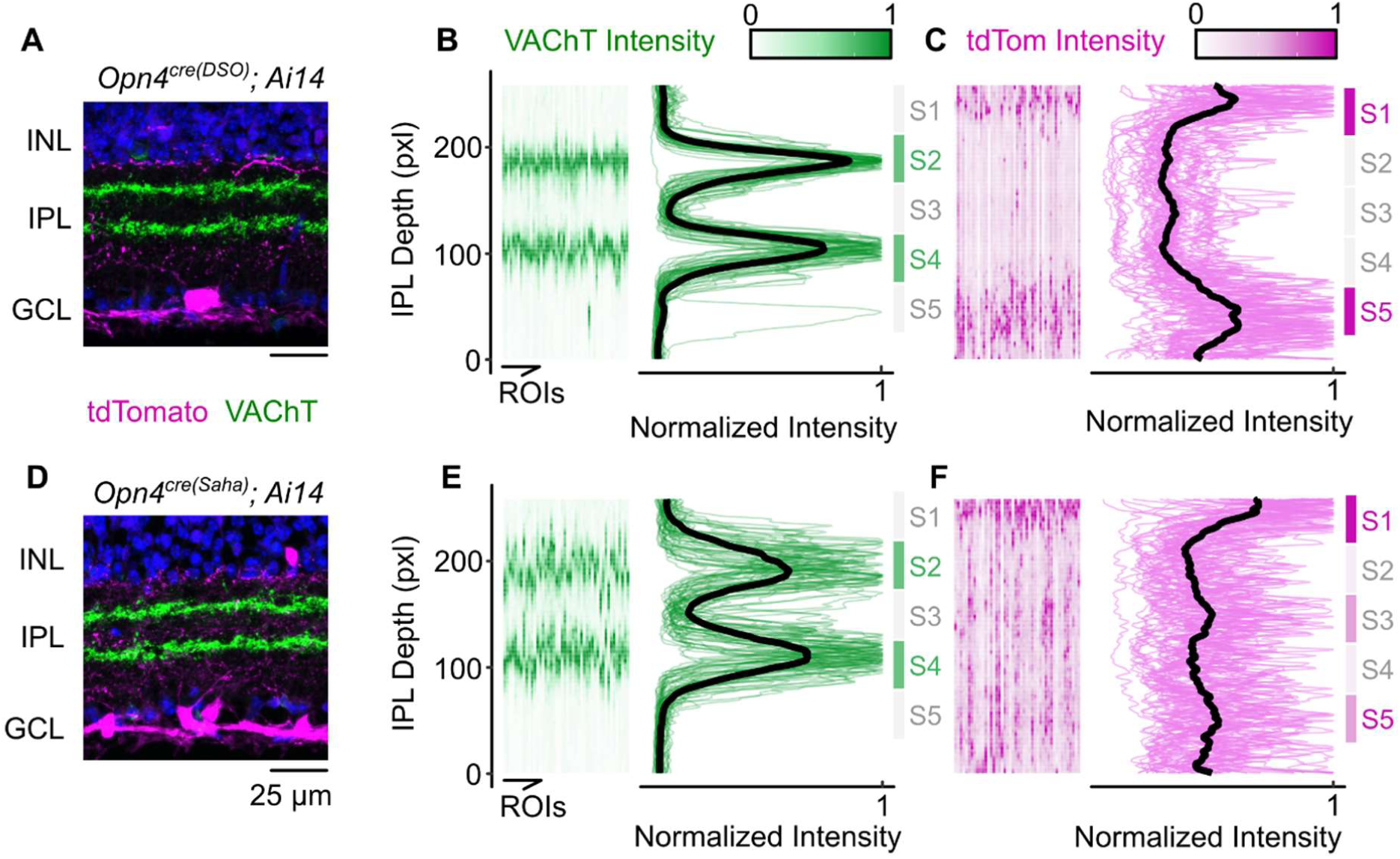
Dendritic lamination within the plexiform layer in both *Cre* lines. (A-C) Analysis of VAChT and tdTomato signal intensity across IPL depth in the *Opn4^cre(DSO^*^)^ line crossed to *Ai14*. VAChT marks starburst amacrine cells (SACs) that laminate in sublamina 2 and 4 (S2 & S4), and is used as an internal control for analysis. (B) Individual 35 x 55 µm ROIs plotted as an intensity heatmap normalized to the maximum intensity value (left) and corresponding average pixel intensity line graph depicting lamination of individual cell types. (D-F) Same as (A-C) but in the *Opn4^cre(Saha^*^)^ line crossed to *Ai14*.

**Supplemental Figure 4.**
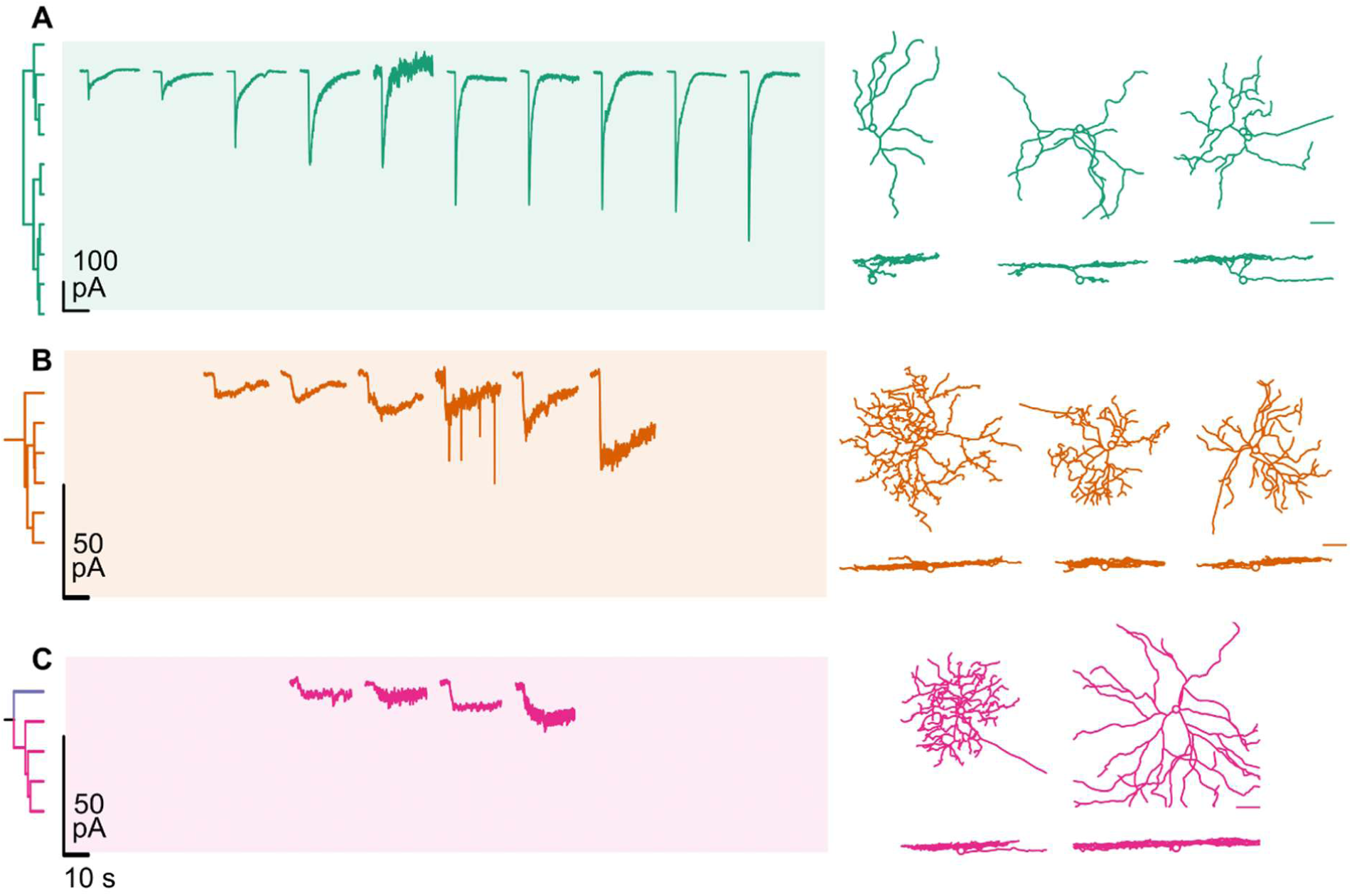
Diversity of intrinsic photocurrents and morphologies of cells in the *Opn4^cre(DSO^*^)^; *Ai9* mouse retina. (A-C) Voltage clamp recordings of intrinsic photocurrents (left) and dye-filled cells (right). Colors and dendrogram reflect clustering of currents highlighted in Figure 4.

**Supplemental Figure 5.**
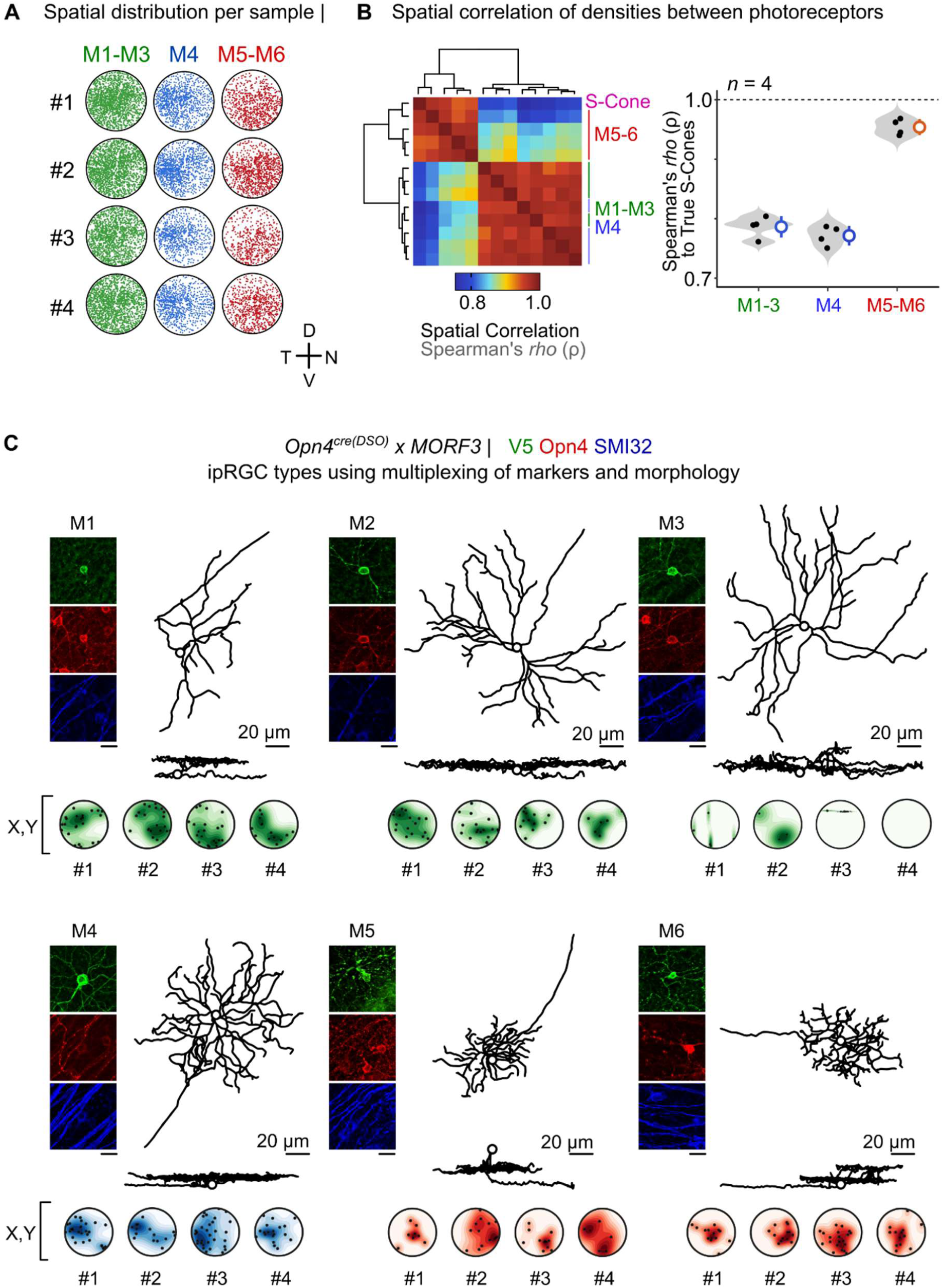
Spatial distributions and single-cell reconstructions with molecular marker multiplexing to identify ipRGC distributions across the retina. (A) Individual polar plots of coarse ipRGC types (M1-M3 tdTomato+ Opn4+; M4 tdTomato+ SMI32+; M5-M6 tdTomato+ Opn4-SMI32-) in the *Opn4^cre(DSO)^; Ai14* line. (B) Spearmann’s correlation of normalized spatial densities of ipRGC types and True S-Cones. Each row and column in the heatmap represent an individual *n* polar plot. (C) Examples of ipRGC types reconstructed with cognate immunofluorescence of V5 (green), Opn4 (red), and SMI32 (blue), with individual polar plots of each type. Number refers to sample ID number.

**Supplemental Figure 6.**
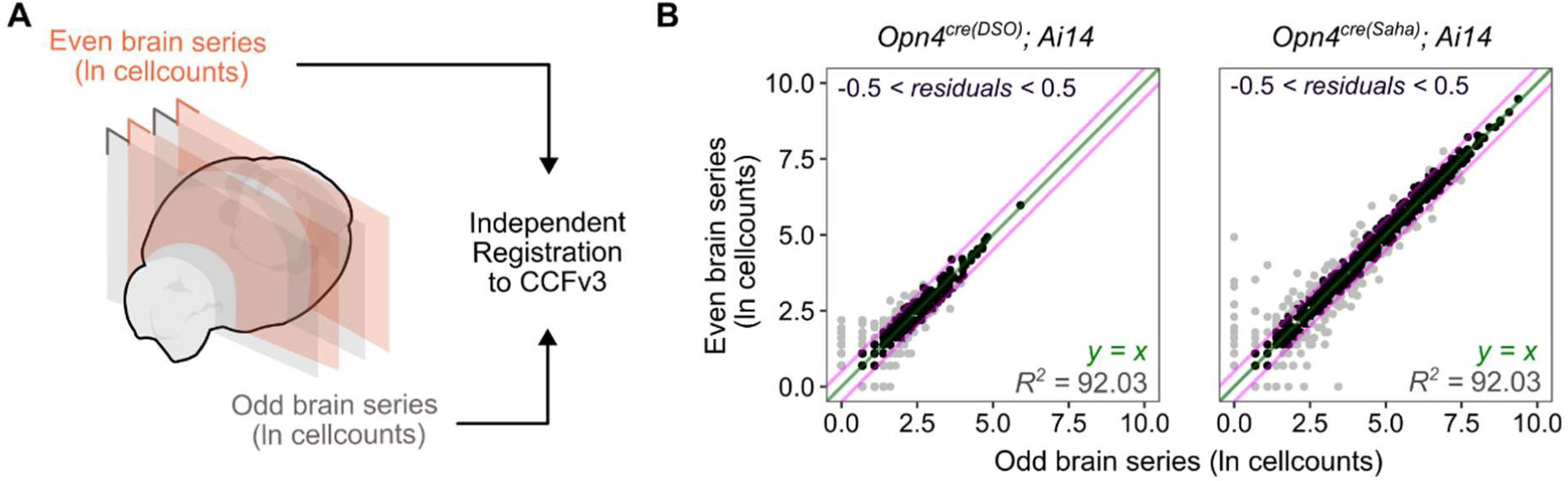
Registration pipeline validation. (A) Schematic representation of registration validation performed on both *Cre* line sections. Series were split into alternating slices (Odd / Even brain series) and independently registered to the Allen CCFv3. (B) Joint linear correlation between odd and even series. Each point represents an individual brain region (∼638), black dots represent residuals between -0.5 and 0.5. Green line represents a linear *y = x* fit, magenta lines represent *y = x +* 0.5 and *y = x* – 0.5.

**Supplemental Figure 7.**
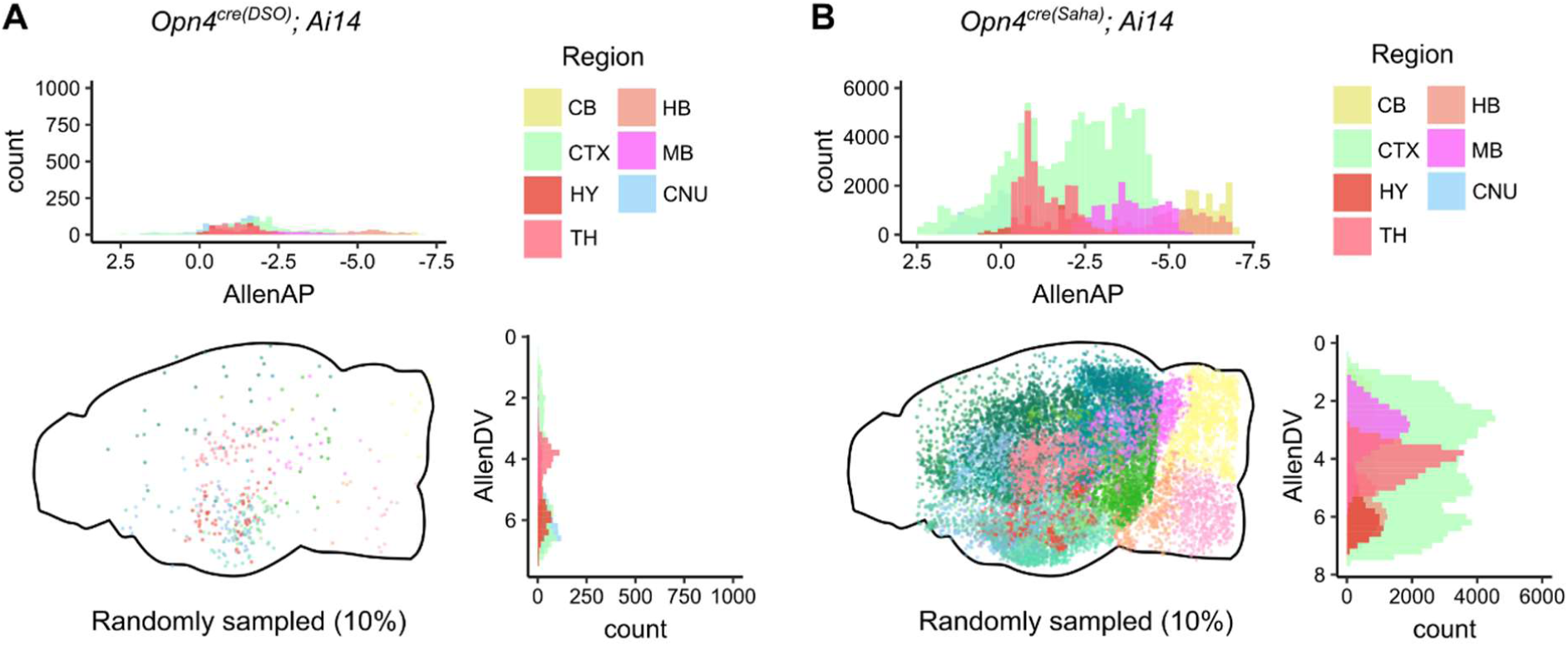
Distribution of cells in the brain of *Opn4cre* mice. (A) Profile view of cell distributions across the *Opn4^cre(DSO^*^)^; *Ai14* brain. (B) Similar to (A) but in the *Opn4^cre(Saha)^; Ai14* line. AllenAP = anterior-posterior axis (mm), AllenDV = dorsal-ventral axis (mm). Notice scale differences in y-axes in count. This was done to highlight cells in the *Opn4^cre(DSO^*^)^ line, as the *Saha* scale would eclipse the histograms in the *DSO* plots.

## REFERENCES

1 Do, M. T. H. Melanopsin and the Intrinsically Photosensitive Retinal Ganglion Cells: Biophysics to Behavior. Neuron 104, 205–226, doi:10.1016/j.neuron.2019.07.016 (2019).

2 Sondereker, K. B., Stabio, M. E. & Renna, J. M. Crosstalk: The diversity of melanopsin ganglion cell types has begun to challenge the canonical divide between image-forming and non-image-forming vision. J Comp Neurol 528, 2044–2067, doi:10.1002/cne.24873 (2020).

3 Schmidt, T. M. & Kofuji, P. Functional and morphological differences among intrinsically photosensitive retinal ganglion cells. J Neurosci 29, 476–482, doi:10.1523/jneurosci.4117-08.2009 (2009).

4 Aranda, M. L. & Schmidt, T. M. Diversity of intrinsically photosensitive retinal ganglion cells: circuits and functions. Cell Mol Life Sci 78, 889–907, doi:10.1007/s00018-020-03641-5 (2021).

5 Berson, D. M., Dunn, F. A. & Takao, M. Phototransduction by retinal ganglion cells that set the circadian clock. Science 295, 1070–1073, doi:10.1126/science.1067262 (2002).

6 Hattar, S., Liao, H. W., Takao, M., Berson, D. M. & Yau, K. W. Melanopsin-containing retinal ganglion cells: architecture, projections, and intrinsic photosensitivity. Science 295, 1065–1070, doi:10.1126/science.1069609 (2002).

7 Göz, D. et al. Targeted destruction of photosensitive retinal ganglion cells with a saporin conjugate alters the effects of light on mouse circadian rhythms. PLoS One 3, e3153, doi:10.1371/journal.pone.0003153 (2008).

8 Hatori, M. et al. Inducible ablation of melanopsin-expressing retinal ganglion cells reveals their central role in non-image forming visual responses. PLoS One 3, e2451, doi:10.1371/journal.pone.0002451 (2008).

9 Güler, A. D. et al. Melanopsin cells are the principal conduits for rod-cone input to non-image-forming vision. Nature 453, 102–105, doi:10.1038/nature06829 (2008).

10 LeGates, T. A. et al. Aberrant light directly impairs mood and learning through melanopsin-expressing neurons. Nature 491, 594–598, doi:10.1038/nature11673 (2012).

11 Fernandez, D. C. et al. Light Affects Mood and Learning through Distinct Retina-Brain Pathways.Cell 175, 71–84.e18, doi:10.1016/j.cell.2018.08.004 (2018).

12 Altimus, C. M. et al. Rods-cones and melanopsin detect light and dark to modulate sleep independent of image formation. Proc Natl Acad Sci U S A 105, 19998–20003, doi:10.1073/pnas.0808312105 (2008).

13 Schmidt, T. M. et al. A role for melanopsin in alpha retinal ganglion cells and contrast detection. Neuron 82, 781–788, doi:10.1016/j.neuron.2014.03.022 (2014).

14 Ecker, J. L. et al. Melanopsin-expressing retinal ganglion-cell photoreceptors: cellular diversity and role in pattern vision. Neuron 67, 49–60, doi:10.1016/j.neuron.2010.05.023 (2010).

15 Chew, K. S. et al. A subset of ipRGCs regulates both maturation of the circadian clock and segregation of retinogeniculate projections in mice. Elife 6, doi:10.7554/eLife.22861 (2017).

16 Schmidt, T. M., Taniguchi, K. & Kofuji, P. Intrinsic and extrinsic light responses in melanopsin-expressing ganglion cells during mouse development. J Neurophysiol 100, 371–384, doi:10.1152/jn.00062.2008 (2008).

17 Chen, S. K., Badea, T. C. & Hattar, S. Photoentrainment and pupillary light reflex are mediated by distinct populations of ipRGCs. Nature 476, 92–95, doi:10.1038/nature10206 (2011).

18 Keenan, W. T. et al. A visual circuit uses complementary mechanisms to support transient and sustained pupil constriction. Elife 5, doi:10.7554/eLife.15392 (2016).

19 Rupp, A. C. et al. Distinct ipRGC subpopulations mediate light’s acute and circadian effects on body temperature and sleep. Elife 8, doi:10.7554/eLife.44358 (2019).

20 Madisen, L. et al. A robust and high-throughput Cre reporting and characterization system for the whole mouse brain. Nat Neurosci 13, 133–140, doi:10.1038/nn.2467 (2010).

21 Maloney, R., Quattrochi, L., Yoon, J., Souza, R. & Berson, D. Efficacy and specificity of melanopsin reporters for retinal ganglion cells. J Comp Neurol 532, e25591, doi:10.1002/cne.25591 (2024).

22 Berg, D. J., Kartheiser, K., Leyrer, M., Saali, A. & Berson, D. M. Transcriptomic Signatures of Postnatal and Adult Intrinsically Photosensitive Ganglion Cells. eNeuro 6, doi:10.1523/eneuro.0022-19.2019 (2019).

23 Vooijs, M., Jonkers, J. & Berns, A. A highly efficient ligand-regulated Cre recombinase mouse line shows that LoxP recombination is position dependent. EMBO Rep 2, 292–297, doi:10.1093/embo-reports/kve064 (2001).

24 Bleckert, A., Schwartz, G. W., Turner, M. H., Rieke, F. & Wong, R. O. Visual space is represented by nonmatching topographies of distinct mouse retinal ganglion cell types. Curr Biol 24, 310–315, doi:10.1016/j.cub.2013.12.020 (2014).

25 Hughes, S., Watson, T. S., Foster, R. G., Peirson, S. N. & Hankins, M. W. Nonuniform distribution and spectral tuning of photosensitive retinal ganglion cells of the mouse retina. Curr Biol 23, 1696–1701, doi:10.1016/j.cub.2013.07.010 (2013).

26 Sonoda, T., Okabe, Y. & Schmidt, T. M. Overlapping morphological and functional properties between M4 and M5 intrinsically photosensitive retinal ganglion cells. J Comp Neurol 528, 1028–1040, doi:10.1002/cne.24806 (2020).

27 Berson, D. M., Castrucci, A. M. & Provencio, I. Morphology and mosaics of melanopsin-expressing retinal ganglion cell types in mice. J Comp Neurol 518, 2405–2422, doi:10.1002/cne.22381 (2010).

28 Duda, S. et al. Spatial distribution and functional integration of displaced ipRGCs. bioRxiv, 2023.2009.2005.556383, doi:10.1101/2023.09.05.556383 (2023).

29 Jain, V., Ravindran, E. & Dhingra, N. K. Differential expression of Brn3 transcription factors in intrinsically photosensitive retinal ganglion cells in mouse. J Comp Neurol 520, 742–755, doi:10.1002/cne.22765 (2012).

30 Valiente-Soriano, F. J. et al. Distribution of melanopsin positive neurons in pigmented and albino mice: evidence for melanopsin interneurons in the mouse retina. Front Neuroanat 8, 131, doi:10.3389/fnana.2014.00131 (2014).

31 Provencio, I. et al. A novel human opsin in the inner retina. J Neurosci 20, 600–605, doi:10.1523/jneurosci.20-02-00600.2000 (2000).

32 Rodriguez, A. R., de Sevilla Müller, L. P. & Brecha, N. C. The RNA binding protein RBPMS is a selective marker of ganglion cells in the mammalian retina. J Comp Neurol 522, 1411–1443, doi:10.1002/cne.23521 (2014).

33 Tran, N. M. et al. Single-Cell Profiles of Retinal Ganglion Cells Differing in Resilience to Injury Reveal Neuroprotective Genes. Neuron 104, 1039–1055.e1012, doi:10.1016/j.neuron.2019.11.006 (2019).

34 Chen, C. K. et al. Characterization of Tbr2-expressing retinal ganglion cells. J Comp Neurol 529, 3513–3532, doi:10.1002/cne.25208 (2021).

35 Lee, E. S., Lee, J. Y., Kim, G. H. & Jeon, C. J. Identification of calretinin-expressing retinal ganglion cells projecting to the mouse superior colliculus. Cell Tissue Res 376, 153–163, doi:10.1007/s00441-018-2964-1 (2019).

36 Do, M. T. H. & Yau, K.-W. Intrinsically Photosensitive Retinal Ganglion Cells. Physiological Reviews 90, 1547–1581, doi:10.1152/physrev.00013.2010 (2010).

37 Schmidt, T. M. & Kofuji, P. Structure and function of bistratified intrinsically photosensitive retinal ganglion cells in the mouse. J Comp Neurol 519, 1492–1504, doi:10.1002/cne.22579 (2011).

38 Estevez, M. E. et al. Form and function of the M4 cell, an intrinsically photosensitive retinal ganglion cell type contributing to geniculocortical vision. J Neurosci 32, 13608–13620, doi:10.1523/jneurosci.1422-12.2012 (2012).

39 Stabio, M. E. et al. The M5 Cell: A Color-Opponent Intrinsically Photosensitive Retinal Ganglion Cell. Neuron 97, 150–163.e154, doi:10.1016/j.neuron.2017.11.030 (2018).

40 Quattrochi, L. E. et al. The M6 cell: A small-field bistratified photosensitive retinal ganglion cell. J Comp Neurol 527, 297–311, doi:10.1002/cne.24556 (2019).

41 Veldman, M. B. et al. Brainwide Genetic Sparse Cell Labeling to Illuminate the Morphology of Neurons and Glia with Cre-Dependent MORF Mice. Neuron 108, 111–127.e116, doi:10.1016/j.neuron.2020.07.019 (2020).

42 D’Souza, S. P. et al. Developmental adaptation of rod photoreceptor number via photoreception in melanopsin (OPN4) retinal ganglion cells. bioRxiv, doi:10.1101/2023.08.24.554675 (2023).

43 Baden, T., Euler, T. & Berens, P. Understanding the retinal basis of vision across species. Nat Rev Neurosci 21, 5–20, doi:10.1038/s41583-019-0242-1 (2020).

44 Wässle, H. & Boycott, B. B. Functional architecture of the mammalian retina. Physiol Rev 71, 447–480, doi:10.1152/physrev.1991.71.2.447 (1991).

45 Berry, M. H. et al. A melanopsin ganglion cell subtype forms a dorsal retinal mosaic projecting to the supraoptic nucleus. Nat Commun 14, 1492, doi:10.1038/s41467-023-36955-6 (2023).

46 Osterhout, J. A. et al. Cadherin-6 mediates axon-target matching in a non-image-forming visual circuit. Neuron 71, 632–639, doi:10.1016/j.neuron.2011.07.006 (2011).

47 Nadal-Nicolás, F. M. et al. True S-cones are concentrated in the ventral mouse retina and wired for color detection in the upper visual field. Elife 9, doi:10.7554/eLife.56840 (2020).

48 Hattar, S. et al. Central projections of melanopsin-expressing retinal ganglion cells in the mouse. J Comp Neurol 497, 326–349, doi:10.1002/cne.20970 (2006).

49 Beier, C., Zhang, Z., Yurgel, M. & Hattar, S. Projections of ipRGCs and conventional RGCs to retinorecipient brain nuclei. J Comp Neurol 529, 1863–1875, doi:10.1002/cne.25061 (2021).

50 Faust, T. E. et al. A comparative analysis of microglial inducible Cre lines. Cell Rep 42, 113031, doi:10.1016/j.celrep.2023.113031 (2023).

51 Bedolla, A. M. et al. A comparative evaluation of the strengths and potential caveats of the microglial inducible CreER mouse models. Cell Rep 43, 113660, doi:10.1016/j.celrep.2023.113660 (2024).

52 Novak, A., Guo, C., Yang, W., Nagy, A. & Lobe, C. G. Z/EG, a double reporter mouse line that expresses enhanced green fluorescent protein upon Cre-mediated excision. Genesis 28, 147–155 (2000).

53 Lobe, C. G. et al. Z/AP, a double reporter for cre-mediated recombination. Dev Biol 208, 281–292, doi:10.1006/dbio.1999.9209 (1999).

54 Sexton, T. J., Bleckert, A., Turner, M. H. & Van Gelder, R. N. Type I intrinsically photosensitive retinal ganglion cells of early post-natal development correspond to the M4 subtype. Neural Dev 10, 17, doi:10.1186/s13064-015-0042-x (2015).

55 Gallego-Ortega, A. et al. Alpha retinal ganglion cells in pigmented mice retina: number and distribution. Front Neuroanat 16, 1054849, doi:10.3389/fnana.2022.1054849 (2022).

56 Behrens, C., Schubert, T., Haverkamp, S., Euler, T. & Berens, P. Connectivity map of bipolar cells and photoreceptors in the mouse retina. Elife 5, doi:10.7554/eLife.20041 (2016).

57 Levine, J. N. & Schwartz, G. W. The Olivary Pretectal Nucleus Receives Visual Input of High Spatial Resolution. bioRxiv, 2020.2006.2023.168054, doi:10.1101/2020.06.23.168054 (2020).

58 Sabbah, S. et al. Intrinsically photosensitive retinal ganglion cells evade temporal filtering to encode environmental light intensity. bioRxiv, 2022.2004.2009.487733, doi:10.1101/2022.04.09.487733 (2022).

59 Andrabi, M., Upton, B. A., Lang, R. A. & Vemaraju, S. An Expanding Role for Nonvisual Opsins in Extraocular Light Sensing Physiology. Annu Rev Vis Sci 9, 245–267, doi:10.1146/annurev-vision-100820-094018 (2023).

60 Zhang, K. X. et al. Violet-light suppression of thermogenesis by opsin 5 hypothalamic neurons. Nature 585, 420–425, doi:10.1038/s41586-020-2683-0 (2020).

61 Bárez-López, S., Bishop, P., Searby, D., Murphy, D. & Greenwood, M. P. Male rat hypothalamic extraretinal photoreceptor Opsin3 is sensitive to osmotic stimuli and light. J Neuroendocrinol 36, e13363, doi:10.1111/jne.13363 (2024).

62 Upton, B. A. et al. Comprehensive Behavioral Analysis of Opsin 3 (Encephalopsin)-Deficient Mice Identifies Role in Modulation of Acoustic Startle Reflex. eNeuro 9, doi:10.1523/eneuro.0202-22.2022 (2022).

63 Matynia, A. et al. Peripheral Sensory Neurons Expressing Melanopsin Respond to Light. Front Neural Circuits 10, 60, doi:10.3389/fncir.2016.00060 (2016).

64 Wang, Q. et al. Synergistic Signaling by Light and Acetylcholine in Mouse Iris Sphincter Muscle. Curr Biol 27, 1791–1800.e1795, doi:10.1016/j.cub.2017.05.022 (2017).

65 Sikka, G. et al. Melanopsin mediates light-dependent relaxation in blood vessels. Proc Natl Acad Sci U S A 111, 17977–17982, doi:10.1073/pnas.1420258111 (2014).

66 Ondrusova, K. et al. Subcutaneous white adipocytes express a light sensitive signaling pathway mediated via a melanopsin/TRPC channel axis. Sci Rep 7, 16332, doi:10.1038/s41598-017-16689-4 (2017).

67 Sonoda, T. et al. A noncanonical inhibitory circuit dampens behavioral sensitivity to light. Science 368, 527–531, doi:10.1126/science.aay3152 (2020).

68 Collins, E. C., Pannell, R., Simpson, E. M., Forster, A. & Rabbitts, T. H. Inter-chromosomal recombination of Mll and Af9 genes mediated by cre-loxP in mouse development. EMBO Rep 1, 127–132, doi:10.1093/embo-reports/kvd021 (2000).

69 Zong, H., Espinosa, J. S., Su, H. H., Muzumdar, M. D. & Luo, L. Mosaic analysis with double markers in mice. Cell 121, 479–492, doi:10.1016/j.cell.2005.02.012 (2005).

70 Liu, J. et al. Non-parallel recombination limits Cre-LoxP-based reporters as precise indicators of conditional genetic manipulation. Genesis 51, 436–442, doi:10.1002/dvg.22384 (2013).

71 Tong, Q. et al. Synaptic glutamate release by ventromedial hypothalamic neurons is part of the neurocircuitry that prevents hypoglycemia. Cell Metab 5, 383–393, doi:10.1016/j.cmet.2007.04.001 (2007).

72 Meng, F. et al. New inducible genetic method reveals critical roles of GABA in the control of feeding and metabolism. Proc Natl Acad Sci U S A 113, 3645–3650, doi:10.1073/pnas.1602049113 (2016).

73 Warren, E. J., Allen, C. N., Brown, R. L. & Robinson, D. W. Intrinsic light responses of retinal ganglion cells projecting to the circadian system. Eur J Neurosci 17, 1727–1735, doi:10.1046/j.1460-9568.2003.02594.x (2003).

74 Walker, M. T., Brown, R. L., Cronin, T. W. & Robinson, P. R. Photochemistry of retinal chromophore in mouse melanopsin. Proc Natl Acad Sci U S A 105, 8861–8865, doi:10.1073/pnas.0711397105 (2008).

75 Lauridsen, K. et al. A Semi-Automated Workflow for Brain Slice Histology Alignment, Registration, and Cell Quantification (SHARCQ). eneuro 9, ENEURO.0483-0421.2022, doi:10.1523/eneuro.0483-21.2022 (2022).

